# Protocol for Magnetic Resonance Imaging Acquisition, Quality Assurance, and Quality Check for the Accelerator Program for Discovery in Brain Disorders using Stem Cells

**DOI:** 10.1101/2020.07.07.191155

**Authors:** Pravesh Parekh, Gaurav Vivek Bhalerao, Rashmi Rao, Vanteemar S. Sreeraj, Bharath Holla, Jitender Saini, Ganesan Venkatasubramanian, John P. John, Sanjeev Jain, ADBS consortium

**Affiliations:** ADBS Neuroimaging Centre, National Institute of Mental Health and Neurosciences (NIMHANS), Bangalore - 560029, India; Multimodal Brain Image Analysis Laboratory, National Institute of Mental Health and Neurosciences (NIMHANS), Bangalore - 560029, India; Translational Psychiatry Lab, National Institute of Mental Health and Neurosciences (NIMHANS), Bangalore - 560029, India; Departments of Psychiatry, National Institute of Mental Health and Neurosciences (NIMHANS), Bangalore - 560029, India; Departments of Neuroimaging and Interventional Radiology, National Institute of Mental Health and Neurosciences (NIMHANS), Bangalore - 560029, India; Departments of Clinical Neurosciences, National Institute of Mental Health and Neurosciences (NIMHANS), Bangalore - 560029, India

**Keywords:** ADBS, Magnetic Resonance Imaging, Longitudinal study, Quality Assurance, Quality Check

## Abstract

**Objective:** The Accelerator Program for Discovery in Brain Disorders using Stem Cells (ADBS) is a longitudinal study focused on collecting and analysing clinical, neuropsychological, neurophysiological, and multimodal neuroimaging data from five cohorts of patients with major psychiatric disorders from genetically high-risk families, their unaffected first-degree relatives, and healthy subjects. Here, we present a complete description of the acquisition of multimodal MRI data along with the quality assurance (QA) and quality check (QC) procedures that we are following in this study.

**Methods:** The QA procedure consists of monitoring of different quantitative measurements using an agar gel and a geometrical phantom. For the already acquired data from human subjects, we describe QC steps for each imaging modality. To quantify reliability of outcome measurements, we perform test-retest reliability on human volunteers.

**Results:** We have presented results from analysis of phantom data and test-retest reliability on a human volunteer. Results show consistency in data acquisition and reliable quantification of different outcome measurements.

**Conclusion:** The acquisition protocol and QA-QC procedures described here can yield consistent and reliable outcome measures. We hope to acquire and eventually release high quality longitudinal neuroimaging dataset that will serve the scientific community and pave the way for interesting discoveries.

## Introduction

Comprehending the neurobiological basis of psychiatric disorders is challenging. There is evidence that suggests various psychiatric disorders have overlapping genetic, environmental, and developmental risk factors (Buckholtz & Meyer-Lindenberg, 2012; Craddock & Owen, 2010; Docherty, Moscati, & Fanous, 2016; Ethridge et al., 2012; McTeague, Goodkind, & Etkin, 2016; McTeague et al., 2017; Meda et al., 2012). This makes it difficult to make specific inferences about the neurobiology of these disorders. Understanding the differential neural basis across disorders may lead to improved insights that may affect diagnosis, prognosis, and treatment planning. New avenues are being explored with advancements in stem cells, genomics, and neurophysiological modes of investigation of brain disorders. While helpful in understanding the disease pathology, cross-sectional studies do not provide any information about the progression of these disorders. A longitudinal study will allow us to monitor the developmental changes over the course of illnesses. Studies like the Human Connectome Project (https://neuroscienceblueprint.nih.gov/human-connectome/connectome-programs) and Alzheimer’s Disease Neuroimaging Initiative (ADNI) (http://adni.loni.usc.edu) have not only led to deeper insights but also generated large databases, thus maximizing the scope of research in clinical neuroscience. A systems-based approach with comprehensive clinical phenotyping would be essential in this process (Saxe et al., 2016). In order to disentangle the shared neurodevelopmental trajectories of these disorders, it is necessary to study samples of patients over time, as well as their unaffected first-degree relatives (FDRs) and socio-demographically matched healthy subjects.

Discovery Biology of Neuropsychiatric Syndromes (DBNS) is an approach that proposes to evaluate overlapping and unique genetic, environmental, and developmental factors across five major psychiatric disorders (schizophrenia, bipolar disorder, obsessive-compulsive disorder, Alzheimer’s dementia and alcohol dependence syndrome) (Viswanath et al., 2018). One of the primary objectives of the Accelerator program for Discovery in Brain disorders using Stem cells (ADBS) project (refer Supplementary Materials) is to create a dataset of patients and their FDRs from families with high genetic loading (with at least another affected FDR with any of the five disorders), and socio-demographically matched healthy subjects. This dataset will consist of detailed clinical and neuropsychological evaluation, stem cells, neurophysiological data, and multimodal neuroimaging data. This will allow us to characterize the structure and function of the brain at a microscopic and macroscopic level across these disorders.

MRI allows us to non-invasively study the structure and functioning of the brain. Precise brain characterization depends on the standards of acquisition and processing of MRI data. Regular monitoring of scanner performance and protocol adherence will ensure reliable outcomes by minimizing subject-independent variances in MRI measurements (Bennett & Miller, 2010; Gunter et al., 2009; Maclaren, Han, Vos, Fischbein, & Bammer, 2014). Given the vast number of sources of noise in MRI signal, ensuring longitudinal quality and reliability is of utmost importance. This can be done in two phases: quality assurance and quality check. Typically, quality assurance (QA) is done at the MRI site by acquiring phantom scans that can help in identifying scanner related artefacts and signal fluctuation, while quality check (QC) is done by manual and/or automated inspection of the already acquired data. The goal of this paper is to provide an overview of the MRI acquisition and QA-QC protocol for the ADBS project. We comprehensively describe each step of acquisition and QA-QC protocol along with example results which will illustrate that the data being acquired under the ADBS project is capable of yielding consistent and reliable outcome.

## Methods

### Participants

The study design, short-term and long-term objectives, and details of sample size calculation of the ADBS project are fully described elsewhere (Viswanath et al., 2018). Participants are recruited after obtaining written informed consent duly approved by institutional ethical committee (Ethics approval number: Item No.VII, Sl.No.7.01., Behavioural Sciences & Item No.XI, Sl.No.11.05., Behavioural Sciences).

### MRI acquisition

MRI acquisition is performed temporally as close as possible to other clinical and endophenotype assessments. All subjects are comprehensively screened for MR safety by the clinical team, followed by on-site screening prior to MRI acquisition (MRI screening form is provided in the Supplementary Materials). Training of subjects for task-based functional MRI (fMRI) is conducted prior to the MRI acquisition session. Primary language (or the language most comfortable among the available languages) of the individual is identified using a screening questionnaire. Task introduction and training is performed by trained neuropsychologists using a computerized independent version of the tasks.

Data is being acquired on a 3T Philips Ingenia CX (Philips Healthcare, Best, The Netherlands) machine using a 32-channel phased-array coil. A fiducial marker (vitamin E capsule) is used for enabling unambiguous delineation of laterality. A plastic mirror is mounted on the head coil such that the participants can view the MR-compatible monitor. Stimulus presentation is done via E-Prime 3 (https://pstnet.com/products/e-prime). Synchronization of fMRI stimulus presentation with the scanner is achieved via NordicNeuroLab (https://www.nordicneurolab.com/en/products) sync box and the stimulus is displayed on an LCD monitor from NordicNeuroLab. An MR-compatible microphone from OptoAcoustic (http://www.optoacoustics.com) with active noise cancellation is used to collect vocal responses from participants.

The MRI acquisition protocols were adapted from the ADNI MRI protocols for Philips Ingenia (software version R5). A brief description of our acquisition protocol is mentioned below while additional details are presented in **Tables S1–S4**. A figure showing the contrast of each type of image is shown in **Figure S1.** Details of post acquisition data integrity check and data storage are mentioned in the supplementary materials.

#### Survey

A localizer sequence (“survey”) is initially run which captures the position of the head by acquiring three slices in sagittal, coronal, and axial views. Since the total duration of acquisition is long (1 hour 07 minutes), we acquire three different survey scans during the session: one at the beginning, second before the first task-based fMRI (tb-fMRI) (~13 min 47 sec into the scan), and a third before the second tb-fMRI (~35 min 02 sec into the scan). An additional survey is done before the diffusion-weighted imaging (DWI) scan (~53 min 50 sec after the beginning of the scan), if there is an indication that the study participant has moved.

#### Structural scans

We acquire a high resolution T1-weighted single-shot 3D turbo field echo (TFE) image at 1×1×1 mm voxel size, in the sagittal orientation. The field of view (FOV) is set to ensure full brain coverage while ensuring that no fold-over happens. An acceleration factor (SENSE) of 2 is applied in the right to left (RL) direction to reduce the overall time taken for this acquisition.

In addition to the T1-weighted scan, we acquire a 3D multishot TFE phase-sensitive inversion recovery (PSIR) image at 1×1×1 mm voxel size, in the sagittal orientation. A SENSE factor of 2 in the anterior-posterior (AP) and 1.75 in the RL direction is applied for reducing the amount of time taken. The PSIR images are reconstructed so that cerebrospinal fluid (CSF) voxels have negative values. This can potentially lead to better contrast between tissue types (Hou, Hasan, Sitton, Wolinsky, & Narayana, 2005; Moran, Kumar, Karstaedt, & Jackels, 1986; Park, Cho, & Cho, 1986).

We also acquire a T2-weighted structural scan at 1×1×1 mm voxel size in the transverse orientation. This is a multishot turbo spin-echo (SE) sequence with fat-suppression using SPIR technique. Apart from using this image for various structural analyses, this image can be also be used for registration and correction of geometrical distortion in DWI.

Additionally, for each study participant, we acquire a FLAIR image to visualise white matter abnormalities.

#### Functional scans

For functional blood oxygenation level-dependent (BOLD) contrast, echo-planar imaging (EPI) method is used. We acquire and discard (at source) five dummy scans to allow magnetisation level to reach a steady-state. During EPI scans, brain areas such as orbitofrontal cortex (OFC) are susceptible to signal loss; to mitigate this, we employ a second-order pencil beam shimming (PBS) covering the entire brain which attempts to homogenize the MR signal in the shim region. Further, EPI scans often lead to geometrical distortion. To correct for potential geometrical distortion, we acquire fieldmap and opposite polarity phase encoding EPI scans (Jezzard & Balaban, 1995; Jezzard & Clare, 1999; Smith et al., 2004).

##### Resting state fMRI

During resting-state fMRI (rsfMRI) scan, study participants are asked to keep their eyes open. Acquisition is carried out without any stimuli. Since previous research has indicated that at least 10 minutes of rsfMRI data is required for reliable characterization of various network properties (though the timing required varied from 10-80 minutes) (Birn et al., 2013; Gordon et al., 2017), we acquire ~10 minutes of rsfMRI data (275 volumes, TR 2.2s).

##### Task-based fMRI

Abnormalities in emotion recognition (Castellano et al., 2015; Daros, Zakzanis, & Rector, 2014; Elferink, van, & Kessels, 2015; Homorogan et al., 2017; Kohler, Walker, Martin, Healey, & Moberg, 2010) and verbal fluency (Cardenas, Kassem, Brotman, Leibenluft, & McMahon, 2016; Clark et al., 2009; Clarke et al., 2016; Liang et al., 2016; Snyder, Kaiser, Warren, & Heller, 2015; Weiner et al., 2015) have been reported in all the disorders being studied in this project. We use tasks that were previously developed at NIMHANS for tb-fMRI acquistions: Tool for Recognition of Emotions in Neuropsychiatric Disorders (TRENDS) (Behere et al., 2008), which is a standardized validated tool for studying emotion recognition deficits in Indian patients, and Verbal Fluency Task (VFT) (John, Halahalli, Vasudev, Jayakumar, & Jain, 2011), which is a semantic category overt word generation task that is implemented as a blocked design. A brief description of these tasks are provided in the supplementary materials.

To prevent any carry over effect of one task to another, task scans are separated by the PSIR scan (described above) which lasts for about 9 minutes. The tasks are further spaced by the preceding two reference scans before the beginning of the actual task (32 seconds each). The tasks are administered in a counterbalanced manner to avoid order effects across participants.

#### Diffusion-weighted imaging

The DWI scan is preceeded by reference images which can be used for correction of EPI-induced geometrical distortions. These are SE images acquired in opposite phase encoding directions (PA and AP), two volumes in each direction. The main scan is acquired with a multi-shell sampling protocol, optimized to provide uniform coverage on each shell, and a global uniform angular coverage (Caruyer, Lenglet, Sapiro, & Deriche, 2013). Three high *b*-value shells corresponding to 1000, 2000, and 3000 (smm^-2^) with 25, 24 and 24 gradient directions respectively and seven interspersed volumes without diffusion weighting (*b*-value = 0) are acquired using a second-order PBS covering whole brain to homogenize the signal in regions susceptible to signal loss.

### Quality Assurance

#### MRI Phantoms

Acquisition of data on MRI phantoms is a way to ensure that the scanner stability is maintained over a period of time. Since there are no sources of variation in a phantom, if acquisition protocol and related factors are kept constant, the scan-to-scan change seen in phantom data should be minimal. Depending on the type of phantom and the data being acquired, a variety of parameters targeting specific scanner or image properties can be studied. In this study, we use two kinds of phantoms: an agar gel phantom for ensuring temporal stability of EPI data, and a geometric phantom for ensuring the stability of acquired MR signal.

##### Agar gel phantom

We use an agar gel phantom (http://pro-project.pl/pro-mri_agar) to quantify the temporal stability of EPI BOLD signal. The phantom is placed as centrally as possible in the 32-channel head coil as shown in **Figure S2** with appropriate paddings. The phantom is placed in the horizontal direction and is corrected for any misalignment based on the localizer scan as shown in **Figure S3**.

EPI BOLD images are acquired on the phantom. The acquisition protocol follows the rsfMRI protocol used for human participants (detailed in **Table S2**) except that only 140 volumes are acquired, and dummy scans are not acquired. The PBS is adjusted so that it covers the entire phantom as shown in **Figure S3.**

To examine the properties of EPI images acquired on the phantom, we followed the methods detailed earlier (Vogelbacher et al., 2018). The implementation is adapted from the MATLAB scripts for the gel phantom implemented in LAB-QA2GO (Vogelbacher et al., 2019). We quantified the signal to noise ratio (SNR), the percent integral uniformity (PIU), percent signal ghosting (PSG), signal to fluctuation noise ratio (SFNR), drift, percent fluctuation, and percent signal change (PSC) for the acquired data (see **Table 1** and supplementary material).

**Table 1:** Summary of quality measures from (Vogelbacher et al., 2018)

| Measure | Significance |
| --- | --- |
| Signal-to-noise ratio (SNR) | Quantification of dominance of meaningful signal over noise in the image |
| Percent integral uniformity (PIU) | Describes uniformity of an image |
| Percent signal ghosting (PSG) | Assessment of the amount of ghosting in the image |
| Signal-to-fluctuation-noise-ratio (SFNR) | Measure of relative temporal noise |
| Percent fluctuation | Assessment of signal variance |
| Drift |  |
| Percent Signal Change | Assessment of homogeneity of SNR |

##### HPD Phantom

The second phantom which we use is the quantitative MRI system phantom from High Precision Devices (HPD) (http://hpd-online.com/system-phantom.php). The phantom is placed as centrally as possible within the head coil and a notch in the mounting plate is used for further alignment. A single coronal slice covering the proton density (PD) array, consisting of 14 spheres with differing proton fractions (5-40 incrementing by 5% and 40-100 incrementing by 10%), is acquired using a SE sequence (slice thickness 6mm, TR=5000ms, TE=10ms, flip angle=90°). Since the PD signal is expected to linearly increase with proton fraction; this property can be used as a proxy for scanner stability.

For each PD image, we detect spheres using the image processing toolbox (MATLAB) and label them using spatial heuristics based on the location of three fiducial points on the slice (see **Figure 1a**). The detected boundary is eroded twice to ensure that only intensities within the sphere are considered. We calculate the mean intensity within the sphere and fit a liner regression model, explaining the mean sphere signal as a function of proton fraction percentages. The percentage variance explained by the model, *R*-squared, is then a summary measure of the linear increase of signal with increasing proton fraction (see **Figure 1**b).

**Figure 1:**
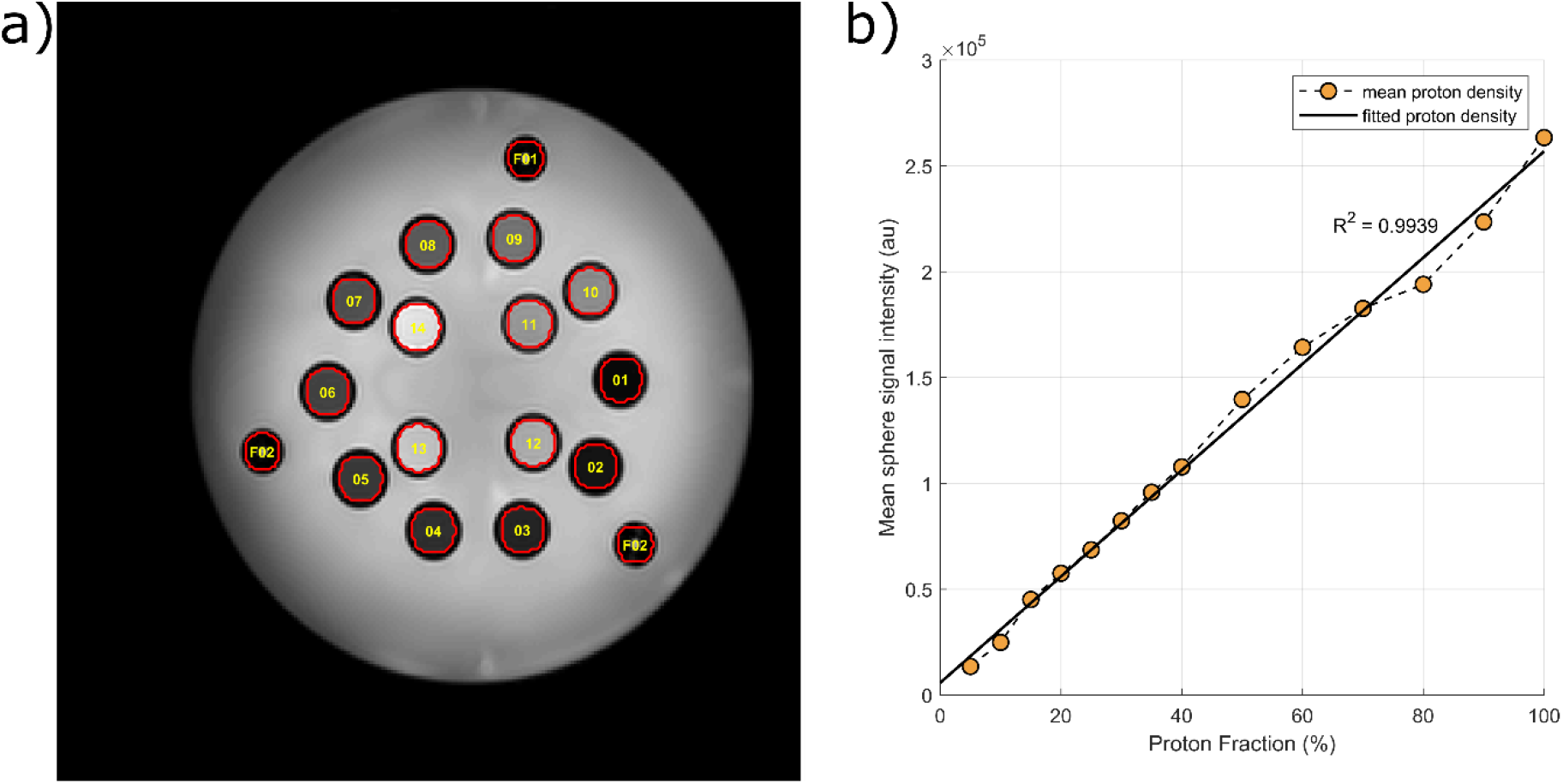
Analysis of proton density images from the HPD phantom; a) fiducials F01, F02, and F03 are first detected. These locations are then used for labeling the other spheres using heuristics. The boundary of the spehres are twice eroded to ensure that only the signal within the sphere is used for any calculation; b) mean signal from each sphere is plotted against the proton fraction in these spheres. Since the signal is supposed to linearly increase with increasing proton fraction, we fit a linear regression model and calculate the *R*-squared value. A poorer fit of this linear model would indicate a problem with the acquisition

#### Test-retest reliability analysis

Unlike phantom data, outcome measures on human data can vary based on factors like (but not limited to) scanner variations, physiological conditions, circadian variations, etc. Establishing the validity of outcome measures is challenging given the lack of ground truth; however, reliability of these measures can be established by repeated measurements on the same subject.

Since we are interested in ensuring reliability of outcome measurements over time, we perform repeated scans on two volunteers who are scanned on three consecutive days approximately in the middle of the month (henceforth referred as acquisition series). This test-retest dataset can be used for quantifying the variation in the assessment of outcome measures (such as brain volumes) each month and by extension, whether these variations are consistent over time. We would expect that even though outcome measures may change over time, the variation seen in them would remain comparable. As proof of concept, we here present reliability analysis on T1-weighted and DWI scans on data from one volunteer (male, right-handed, 29 years old) for the time between July 2019 – December 2019 (three acquisitions per month, 18 total scans; see **Table S7** for scan schedule).

Between July and September, the volunteer was on oral medication for a medical condition; medication details are mentioned in **Table S8**.

##### Reliability of T1-weighted scans

To quantify the reliability of T1-weighted scans, we perform segmentation of T1-weighted images using SPM12 (v7487) (https://www.fil.ion.ucl.ac.uk/spm/) and calculate total gray matter (GM), white matter (WM), CSF, and total intracranial volumes (TIV). For each acquisition series, for each measurement, we calculate the percentage coefficient of variation (CV), defined as the ratio of standard deviation of tissue volume to the mean tissue volume.

For this proof of concept analysis, we tested the hypothesis that the mean CV for any volumetric measurement between July-September (series1, 9 scans) was the same as the mean CV for the same volumetric measurement between October-December (series2, 9 scans). We used permutation testing to test this hypothesis. First, we quantified the absolute difference in CVs between the two series (*CV_diff_*). Then, we randomized the series labels. The maximum number of ways that this can be done is 48,620 (including the original series membership). For each of these arrangements of permuted series labels, we computed the new CV. The two-sided *p*-value was calculated as the fraction of times the absolute difference in permuted CV became equal to or exceeded the originally observed difference *CV_diff_*.

##### Segmentation consistency

For the same volunteer, we also calculated pairwise Dice coefficient (DC) to assess the consistency of segmentation for each of the three tissue classes. We binarized the GM, WM, and CSF segmentation using a liberal threshold of 0.1; then for each pair of images (18 total images, 153 pairwise combinations), we calculated DC for each tissue class. Since the pairs will consist of time points within and across months, this will highlight the consistency of segmentation within shorter and over longer periods of time.

##### Reliability of diffusion scans

Similar to the procedure described for T1-weighted scans, to examine the reliability of diffusion scans, we split the data into two series: series1 (July-September, 9 scans) and series2 (October-December, 9 scans). For each diffusion scan, using FSL 6.0.1, we performed distortion, eddy current, and motion correction as described in Quality Check section (see below). We fit a weighted least squares (WLS) tensor model and obtained fractional anisotropy (FA), radial diffusivity (RD), axial diffusivity (AD), and mean diffusivity (MD) maps for each image. We then used tract-based spatial statistics (TBSS) pipeline to obtain a skeleton image which was used for calculating the whole brain average FA, RD, AD, and MD values for each scan. We followed the same permutation testing method as that for T1-weighted scans.

#### Change in quality assurance schedule

From July 2019, we acquire EPI data on agar gel phantom daily while the PD acquisition on HPD phantom happens on a weekly basis. This explains the increased number of time points from July onwards in the results of these phantoms (**Figure 2** and **Figure 3**). This increase in the number of phantom scans can help us better monitor scanner performance and ensure high quality data acquisition.

**Figure 2:**
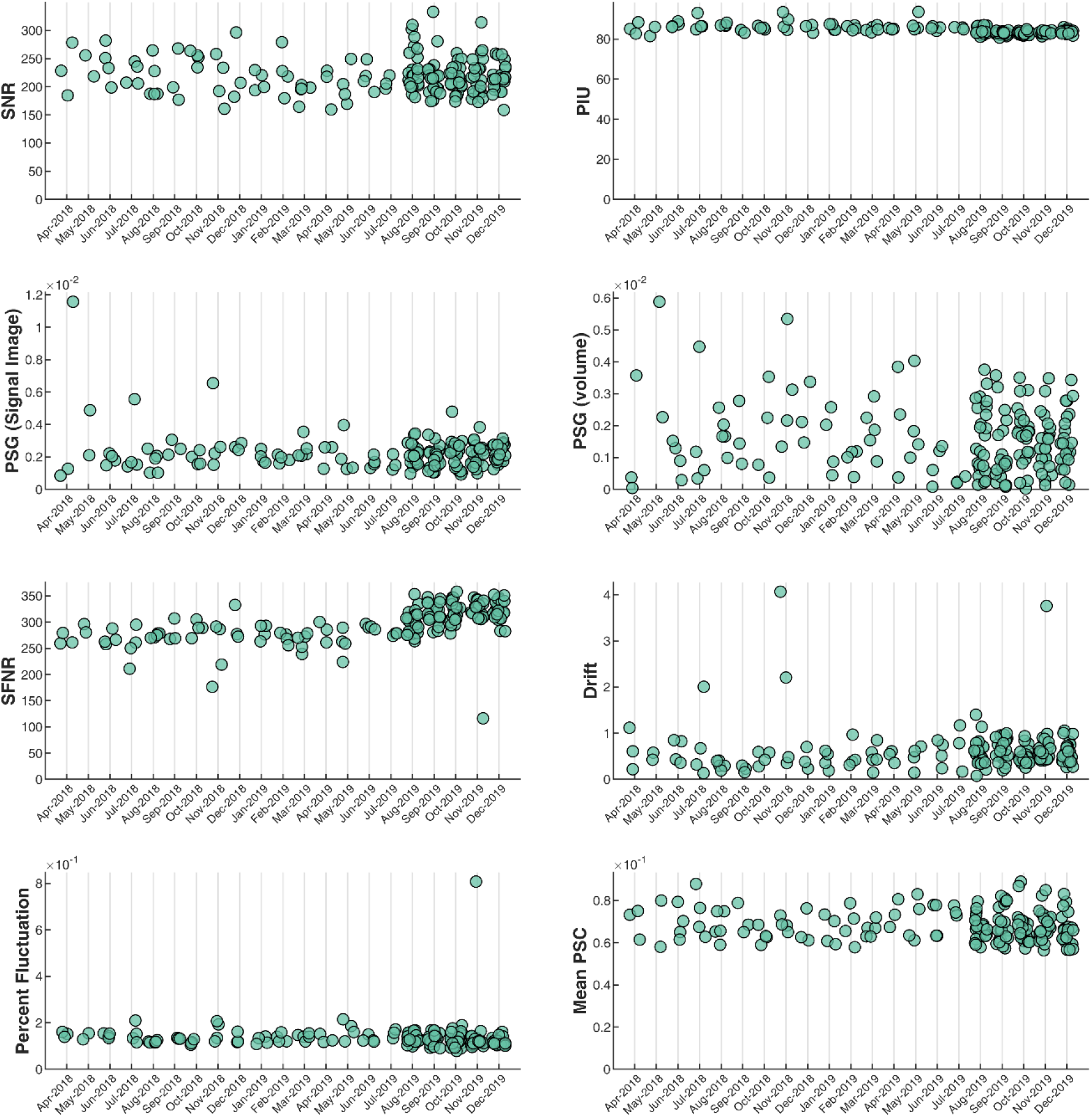
Variation of EPI quality measures on agar gel phantom (see **Table S5** for a breakup of number of scans per month); SNR: signal-to-noise ratio, PIU: percent integral uniformity, PSG: percent signal ghosting, SFNR: signal-to-fluctuation-noise-ratio, PSC: percent signal change

**Figure 3:**
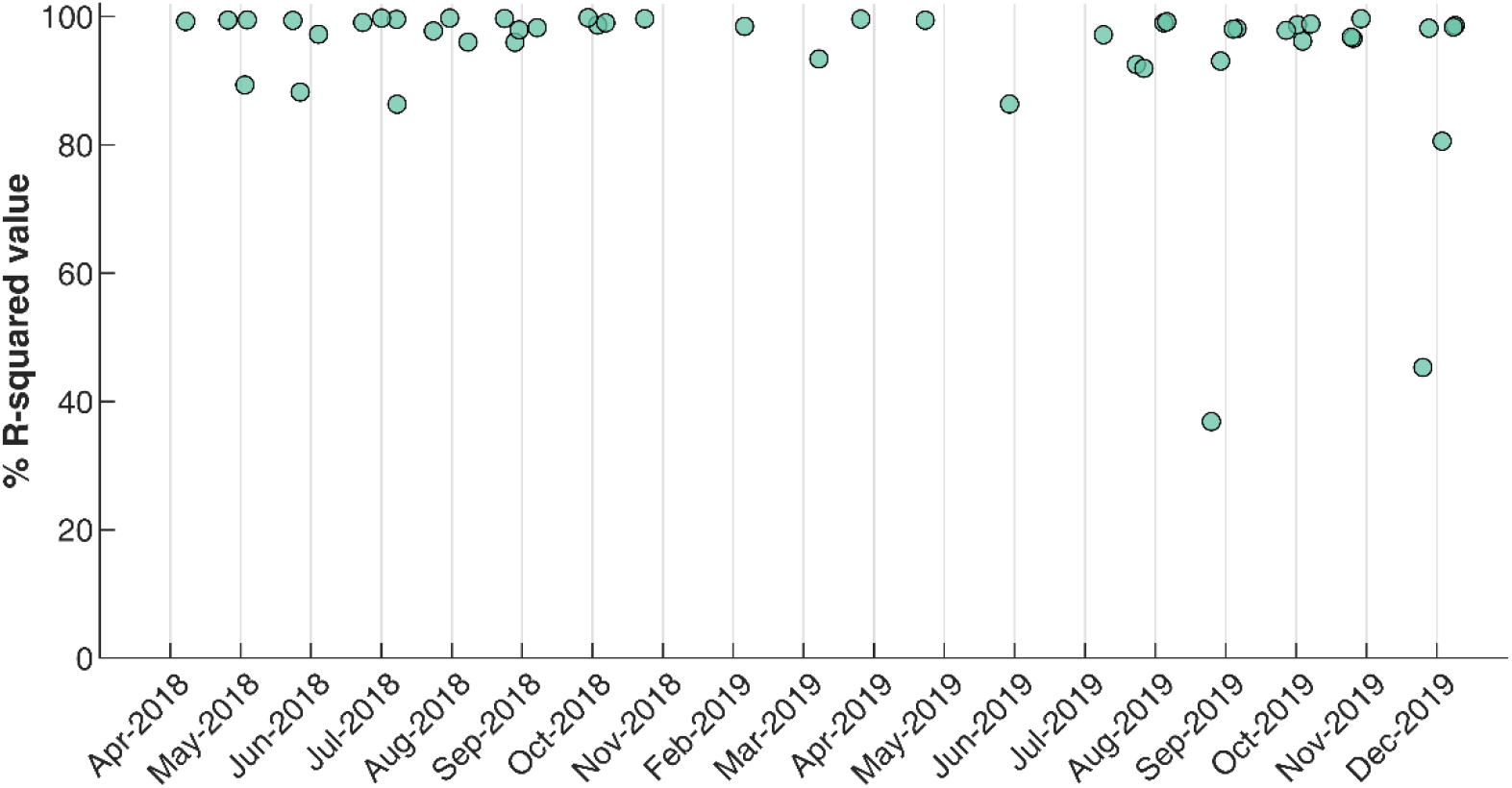
Percentage *R*-squared value from proton density images acquired on HPD phantom between April 2018 and December 2019 (see **Table S6** for a breakup of number of scans per month)

### Quality Check

#### Structural pipeline

Each structural image is reviewed by a neuroradiologist and a report is generated. If any significant structural abnormality is found, the study participant is flagged for further clinical evaluation. Following this, we set the origin of the images to the anterior commissure (AC). This step is done using acpcdetectv2 (part of Automatic Registration Toolbox) (https://www.nitrc.org/projects/art). In case this step fails, we manually set the origin using SPM12 (https://www.fil.ion.ucl.ac.uk/spm/).

##### CAT and MRIQC

All T1-weighted images (with origin set to the AC) are segmented using the Computational Anatomy Toolbox (CAT) (http://www.neuro.uni-jena.de/cat/). The quality check module of CAT returns an image quality grade (A+, A, A-, to F). Any image receiving a rating of “C” or below is flagged. We also use automated quality prediction and visual reporting of MRI scans by running MRIQC (Esteban et al., 2017) on structural scans which generates a detailed visual and quantitative report for each image. Additionally, we use the pre-trained random forest classifier of MRIQC to predict if a T1-weighted image is of good quality or not.

#### Functional pipeline

##### Minimal preprocessing

The minimal preprocessing pipeline for functional images consists of motion correction (using SPM’s realign and unwarp). We then perform slice timing correction and apply the transforms from the T1 AC-PC detection step to the functional image, followed by co-registering the functional image to AC-PC aligned T1-weighted image. The T1-weighted scan is segmented, and a deformation field is estimated which non-linearly warps the brain from native space to the MNI space. This forward deformation field is applied to the co-registered functional scans to warp functional images to the MNI space. The bounding box during normalization step is calculated to allow full brain coverage (rather than using SPM’s default bounding box).

##### Motion profile

At each stage of preprocessing of functional data, we estimate the number of time points that can be considered as motion outliers. This is done using a modification of the DVARS (*D* referring to temporal derivative of time courses, VARS referring to root-mean-squared variance over voxels) approach (Afyouni & Nichols, 2018). A DVARS *p*-value is computed which can be used for declaring a pair of time points as outliers. Additionally, a liberal user-defined threshold is also applied to detect outliers. An example plot is shown in **Figure S5**.

##### Brainmask profile

Brain regions such as the OFC are susceptible to signal loss. To quantify signal loss, we eliminate all voxels which have a mean signal less than 80% of global signal (GS) (similar to an approach used by SPM). We create a binary mask of the voxels which survive this threshold and examine them visually to identify subjects in which the mask is of poor quality. Additionally, we calculate the ratio of voxels remaining in the mask and the template mask (shipped with SPM). Subjects with drastically reduced proportion of voxels can then be flagged using a box-plot method. An example of these brain mask profiles are shown in **Figure S6b**.

##### Signal profile and left-right correlation

To flag images with severe signal loss, we derive mean time series from regions that are susceptible to signal loss, regions not susceptible to signal loss, GS from left hemisphere, right hemisphere, and average GS. These regions are defined using the Hammers atlas (Faillenot, Heckemann, Frot, & Hammers, 2017; Gousias et al., 2008; Hammers et al., 2003) and comprise of bilateral orbitofrontal gyri, gyrus rectus, superior temporal gyrus, and the lateral occipital lobe. The mean percentage difference can be calculated from each region with respect to the GS. Any subject with a significant reduction in signal with this reference signal can be flagged and eliminated. Additionally, looking at GS independently from left and right hemispheres can be useful in diagnosing any hemispheric signal abnormalities which might happen during acquisition (such as a bad channel in the head coil). We further quantify the correlation between left and right hemispheres and outliers are detected using a box-plot method. An example of this signal profile is shown in **Figure S6a**.

##### MRIQC

Apart from the above-mentioned methods of assessing the quality of functional images, we also run MRIQC for all functional images.

#### Diffusion pipeline

##### Distortion correction

We correct EPI induced distortions by using opposite phase encoding directions reference scans. These non-diffusion scans are passed as an input to FSL’s topup to estimate susceptibility-induced off-resonance field using a method similar to that described in (Andersson, Skare, & Ashburner, 2003) as implemented in FSL (Smith et al., 2004) and the two images are combined into a single corrected one. Once the images are distortion corrected, we derive a binary brain mask using the FSL brain extraction toolbox (BET).

##### Eddy current and motion correction

We employ FSL’s eddy (Andersson & Sotiropoulos, 2016) to simultaneously correct for eddy current and motion. The previously calculated brain mask is passed to eddy along with information on which direction distortion is expected, the *b*-vector and *b*-value files, and the estimated field from topup. Additionally, we use the “repol” option (Andersson, Graham, Zsoldos, & Sotiropoulos, 2016) to replace volumes or slices with partial or complete signal loss (due to movement) with their Gaussian Process predictions. Finally, we also use the “mporder” option (Andersson et al., 2017) to find and replace those slices (within a volume) which are corrupted by motion.

##### Subject and group level quality assessment

Once the data is distortion-, eddy current-, and motion-corrected, we use FSL’s eddy’s QUAD (Bastiani et al., 2019) module for quality assessment. This subject level visual and quantitative report that is generated, derives QC measures based on volume-to-volume motion, within-volume motion, eddy current-induced distortion, susceptibility induced distortion, and the number of outliers replaced. Signal-to-noise and contrast-to-noise ratios are additionally calculated and incorporated in the subject level report. Once QC reports at subject level are ready, group-level reports are generated/updated by using FSL’s eddy’s SQUAD tool.

##### Tensor fitting

Once the data is distortion-, eddy current-, and motion-corrected (and the *b*-vectors rotated), we fit a WLS tensor model to the data using FSL dtifit. Voxel-wise sum of squared errors is also saved, and the principle diffusion direction is visualized by overlaying on the FA map.

## Results

### Phantom QA

#### Agar gel phantom

For agar gel phantom, we calculated SNR, PIU, PSG, SFNR, drift, percent fluctuation, and PSC, which quantify the temporal stability of the EPI images. Descriptive statistics of these quality measures for 172 time points collected between April 2018 and December 2019 is presented in **Table 2** and **Figure 2** shows the variation in these over time.

**Table 2:** Descriptive summary of EPI quality check on agar gel phantom (see text for details)

| **Measure** | **Mean** | **Standard Deviation** | **Min** | **Max** |
| --- | --- | --- | --- | --- |
| **SNR** | 219.4163 | 31.6091 | 158.8507 | 333.1398 |
| **Mean PIU2** | 84.2961 | 2.0638 | 80.7046 | 93.5360 |
| **PSG<sub>signalimage</sub><sup>1</sup>** | 0.0022 | 0.0011 | 0.0008 | 0.0116 |
| **PSG<sub>volume</sub><sup>2</sup>** | 0.0016 | 0.0011 | 0.0000 | 0.0059 |
| **SFNR** | 299.6195 | 33.0544 | 116.2972 | 358.1444 |
| **Drift** | 0.6043 | 0.4615 | 0.0753 | 4.0682 |
| **Percent Fluctuation** | 0.1335 | 0.0570 | 0.0786 | 0.8075 |
| **Mean PSC** | 0.0680 | 0.0071 | 0.0564 | 0.0890 |

<sup>1</sup> Slice (mean\_PSG); <sup>2</sup> Volume (PSG2)

#### HPD phantom

For HPD phantom (PD images), we calculated the percentage *R*-squared value for each time point and examined this value over time. We have shown the fitted percentage *R*-squared value for 48 time points between April 2018 and December 2019 in **Figure 3**. We noticed two clear outliers, one in September 2019 and one in December 2019. Empirical assessment tells us that this was due to an incorrect slice being selected during the acquisition procedure.

### Test-retest reliability analysis

#### Reliability of T1-weighted scans

We used permutation test to test the hypothesis that the whole brain CV of total GM, WM, CSF, and TIV were not statistically different between July-September (series1) and October-December (series2). Summary of these results are shown in **Table 3**.

**Table 3:** Coefficient of variation (CV) and *p*-values for whole brain volumetric measurements from T1-weighted scans for test-retest reliability; series1 refers to 9 scans on the same subject between July-September 2019 and series2 refers to 9 scans on the same subject between October-December 2019 (three scans per month) (see Table S7 for scan schedule)

| Measurement | CV series1 (%) | CV series2 (%) | \| CV<sub>diff</sub> (%) \| | $p$-value |
| --- | --- | --- | --- | --- |
| Total GM volumes | 0.5964 | 0.8234 | 0.2270 | 0.4272 |
| Total WM volumes | 0.5815 | 0.5698 | 0.0118 | 0.9210 |
| Total CSF volumes | 1.0910 | 1.5059 | 0.4149 | 0.5917 |
| TIV | 0.1032 | 0.2101 | 0.1069 | 0.0564 |

#### Consistency of segmentation

Pairwise DC for GM, WM and CSF tissues were calculated to examine the consistency in T1-weighted image segmentation across timepoints. There were total 153 pairs for the volunteer data acquired from July 2019 to December 2019, i.e. 3 acquisitions per month. The graph in **Figure 4** shows that DC for all the tissue segmentations were well above 0.9 for every pair of timepoints.

**Figure 4:**
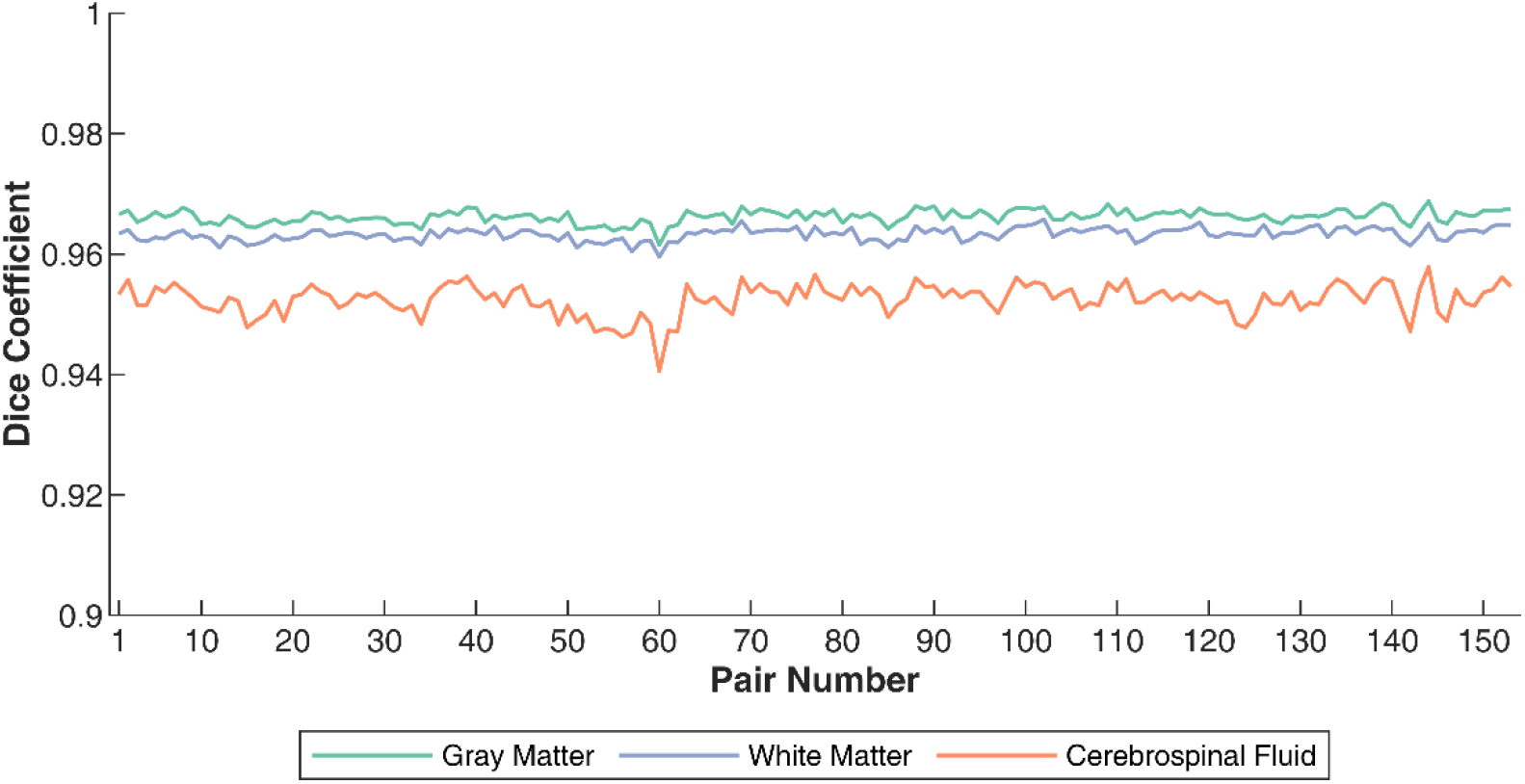
Pariwise Dice coefficient for gray matter (green colour), white matter (purple line), and cerebrospinal fluid (orange colour); between July and December 2019, a volunteer underwent three scans per month (total 18 scans); Dice coefficient between the segmentation of the three tissue classes was calculated for all pairs of images (total 153 pairs) to examine the consistency of segmentation (see **Table S7** for details of reliability schedule)

#### Reliability of diffusion scans

We used permutation testing to test the hypothesis that the whole brain average diffusion measures (FA, AD, RD, and MD) were not statistically different between July-September (series1) and October-December (series2). Summary of these results are shown in **Table 4**. None of the CV of any of the measurements were significantly different between the two assessment series.

**Table 4:** Coefficient of variation (CV) and *p*-values for whole brain average diffusivity measurements from diffusion weighted scans for test-retest reliability; series1 refers to 9 scans on the same subject between July-September 2019 and series2 refers to 9 scans on the same subject between October-December 2019 (three scans per month) (see Table S7 for scan schedule)

| Measurement | CV series1 (%) | CV series2 (%) | \| CV<sub>diff</sub> (%) \| | $p$-value |
| --- | --- | --- | --- | --- |
| Fractional anisotropy | 0.5388 | 0.4224 | 0.1164 | 0.4485 |
| Axial diffusivity | 0.4988 | 0.3220 | 0.1768 | 0.1186 |
| Radial diffusivity | 0.8355 | 0.5157 | 0.3198 | 0.1132 |
| Mean diffusivity | 0.6529 | 0.3824 | 0.2705 | 0.0798 |

## Discussion

Quality assurance and quality check protocols are critical to ensure stable and reliable MRI measurements. Different QA measurements have been reported like SNR and SFNR (Friedman & Glover, 2006), PSC (Stöcker et al., 2005), etc. These measurements can be used to quantify the signal quality of MRI phantoms or human data. Another method of assessing quality of MRI data is by examining the reliability of outcome measures (for example, (Maclaren et al., 2014) examined CV of regional brain volumes over data from three subjects; (Madan & Kensinger, 2017) examined reliability of volumes and surface features using two open-access datasets; (McGuire et al., 2017) worked on three assessments for 25 subjects using multimodal brain imaging data). These approaches show that tracking of outcome variables over repeated assessments can not only highlight the natural biological variability but can also serve as a measure for consistent scanner performance.

In our approach to reliability analysis, to quantify variation on a short-term and a long-term basis, the same subjects are scanned repeatedly. Our test-retest reliability results on structural and diffusion images show that volumetric and diffusivity measurements are reliable over a period of time. Examination of DC shows that segmentations have very high consistency over the same period. Apart from these, visual inspection of the quality measures from both the phantoms show reasonable consistency over time. However, there are some time points which stand out in comparison to the rest of the data. This could be due to a number of factors like handling of the phantom and its positioning, timing of the day, software or hardware changes, or scanner malfluctuation (Vogelbacher et al., 2018). Given differences in scanners, acquisition protocol, types of phantom, and other such reasons, it is not straight forward to establish any range of these QA measurements. (Vogelbacher et al., 2018) have shown a strong dependence on the positioning of the phantom and QA measurements and suggest that the only way to alleviate this is by using a phantom holder. The use of a phantom holder at our site will ensure that the same slice of the phantom is scanned everytime. Additionally, we have plans to integrate an automated warning system to make sure that scanner instabilities can be appropriately detected. This is similar to the idea of LAB-QA2GO (Vogelbacher et al., 2019) but with the modifications relevant to our site and phantom incorporated. Taken together, these show good stabaility of the MR signal as well as stability of structural, functional, and diffusion scans that are being acquired as part of the ADBS project.

Given that ADBS project aims to acquire longitudinal neuroimaging data, implementation of rigorous QA-QC is essential to minimize scanner related variations. Though it is impossible to control for all forms of variations (such as hardware and software changes), monitoring QA measures can help in identifying potential problems. The procedures mentioned here are the QA-QC method being employed during data acquisition which may not necessarily translate to methods of choice for data analyses. Since the ADBS data acquisition relies on a single site/scanner, we do not face inter-site/inter-scanner variability as a factor (unlike multi-site studies). This also allows for a stricter control on variability caused by human factors (for example planning of an acquisition), time window during which acquisition happen, and the incorporation of methods to ensure scanner stability. One of the key aspects that we have focused here is ensuring scanner stability by using different MRI phantoms and incorporating test-retest reliability on human volunteers. As the study progresses, it will be necessary to update some of these QA-QC measures to incorporate the longitudinal nature of the dataset. These methods would also need to be periodically revised as newer methods get proposed and adapted in the field. As of now, we have adapted current methods in the field which require little to no manual intervention. With the influx of data, it will be possible to develop/incorporate machine learning methods for classifying problematic images (we note that the (Alfaro-Almagro et al., 2018) have used a similar idea for identifying problematic T1-weighted images).

Here we here presented ADBS MRI acquisition and QA-QC methods. We hope to curate a high quality neuroimaging database that can serve the scientific community and contribute to understanding the neurobiology of various psychiatric disorders. We also expect that the pipeline for rigorous quality control and reliability measures that are detailed in this manuscript would benefit researchers while planning similar longitudinal neuroimaging studies in various neuropsychiatric disorders.

## Accelerator Program for Discovery in Brain Disorders using Stem cells (ADBS) Consortium

Biju Viswanath^1^, Naren P. Rao^1^, Janardhanan C. Narayanaswamy^1^, Palanimuthu T. Sivakumar^1^, Arun Kandasamy^1^, Muralidharan Kesavan^1^, Urvakhsh Meherwan Mehta^1^, Odity Mukherjee^2^, Meera Purushottam^1^, Ramakrishnan Kannan^1^, Bhupesh Mehta^1^, Thennarasu Kandavel^1^, B. Binukumar^1^, Deepak Jayarajan^1^, A. Shyamsundar^1^, Sydney Moirangthem^1^, K. G. Vijay Kumar^1^, Jayant Mahadevan^1^, Jagadisha Thirthalli^1^, Prabha S. Chandra^1^, Bangalore N. Gangadhar^1^, Pratima Murthy^1^, Mitradas M. Panicker^3^, Upinder S. Bhalla^3^, Sumantra Chattarji^3^, Vivek Benegal^1^, Mathew Varghese^1^, Janardhan Y. C. Reddy^1^, Padinjat Raghu^3^ & Mahendra Rao^3^

^1^National Institute of Mental Health and Neuro Sciences (NIMHANS) institute for Stem Cell Biology and Regenerative Medicine (InStem)

^3^National Center for Biological Sciences (NCBS)

## Acknowledgements

This work is supported by the Department of Biotechnology (DBT), Government of India [BT/PR17316/MED/31/326/2015]; phantom scan data presented in this article were acquired as part of two funded projects: BT/PR17316/MED/31/326/2015 (DBT, Governemnt of India) and RO1 MH113250 (National Institute of Mental Health in the United States). We thank the MRI technicians, Ms. Manasa Venkat and Mr. Vinod Kumar for their assistance in setting up the acquisition protocol and for acquisition of MRI data; and Mr. Bopanna Sugandh, and Mr. Anand Yadav for acquisition of MRI data. We thank T. Nichols and S. Afyouni for their inputs on the usage of the DVARS toolbox. We are grateful to Christoph Vogelbacher and his team for their inputs on phantom analysis. Colour scheme for the figures are from ColorBrewer 2.0 (http://colorbrewer2.org/) by Cynthia A. Brewer, Geography, Pennsylvania State University (accessed 25-Oct-2019). Some of the figures use the tight_subplot function (Pekka Kumpulainen (2020). tight_subplot(Nh, Nw, gap, marg_h, marg_w) (https://www.mathworks.com/matlabcentral/fileexchange/27991-tight_subplot-nh-nw-gap-marg_h-marg_w), MATLAB Central File Exchange. Retrieved January 27, 2020).

## Author’s contributions

**PP**: Conceptualization, Methodology, Software, Formal analysis, Data Curation, Writing - Original Draft, Writing - Review & Editing, Visualization; **GVB**: Conceptualization, Methodology, Software, Formal analysis, Data Curation, Writing - Original Draft, Writing - Review & Editing, Visualization; **RR**: Methodology, Investigation, Data Curation, Writing - Review & Editing; **VSS:** Conceptualization, Investigation, Writing - Original Draft, Writing - Review & Editing, Project administration; **BH:** Conceptualization, Investigation, Writing - Original Draft, Writing - Review & Editing, Project administration; **JS:** Methodology, Supervision, Project administration, Writing - Review & Editing; **GV:** Conceptualization, Methodology, Validation, Resources, Writing - Review & Editing, Supervision, Project administration, Funding acquisition; **JPJ**: Conceptualization, Methodology, Validation, Resources, Writing - Review & Editing, Supervision, Project administration, Funding acquisition; **SJ:** Conceptualization, Methodology, Resources, Writing - Review & Editing, Supervision, Project administration, Funding acquisition; All authors read and approved the final manuscript

## Supplementary Material

### Brief description of the ADBS project

The Discovery Biology of Neuropsychiatric Syndromes (DBNS) is an approach that examines overlapping and unique genetic, environmental, and developmental factors in schizophrenia, bipolar disorder, obsessive-compulsive disorder, Alzheimer’s dementia and alcohol dependence syndrome (Viswanath et al., 2018). Using this approach, the Accelerator program for Discovery in Brain disorders using Stem cells (ADBS) is an ongoing project that aims to recruit 4,500 individuals who undergo phenotypic assessments and blood sampling (for molecular genetic studies and cellular modelling of diseases). Out of these, a subset of 1500 consenting individuals (1,200 from affected families and 300 healthy controls from unaffected families) form the neurodevelopmental endophenotype cohort that is being assessed biennially using structural, functional, and diffusion MRI, electroencephalography (EEG), functional near infra-red spectroscopy (fNIRS), eye movement tracking and neuropsychological evaluation.

## Acquisition protocol

### Structural scans

A summary of critical acquisition parameters for T1w, T1w-PSIR, and T2w scans are presented in **Table S1**. **Figure S1** shows the contrasts of the three structural scans.

**Figure S1:**
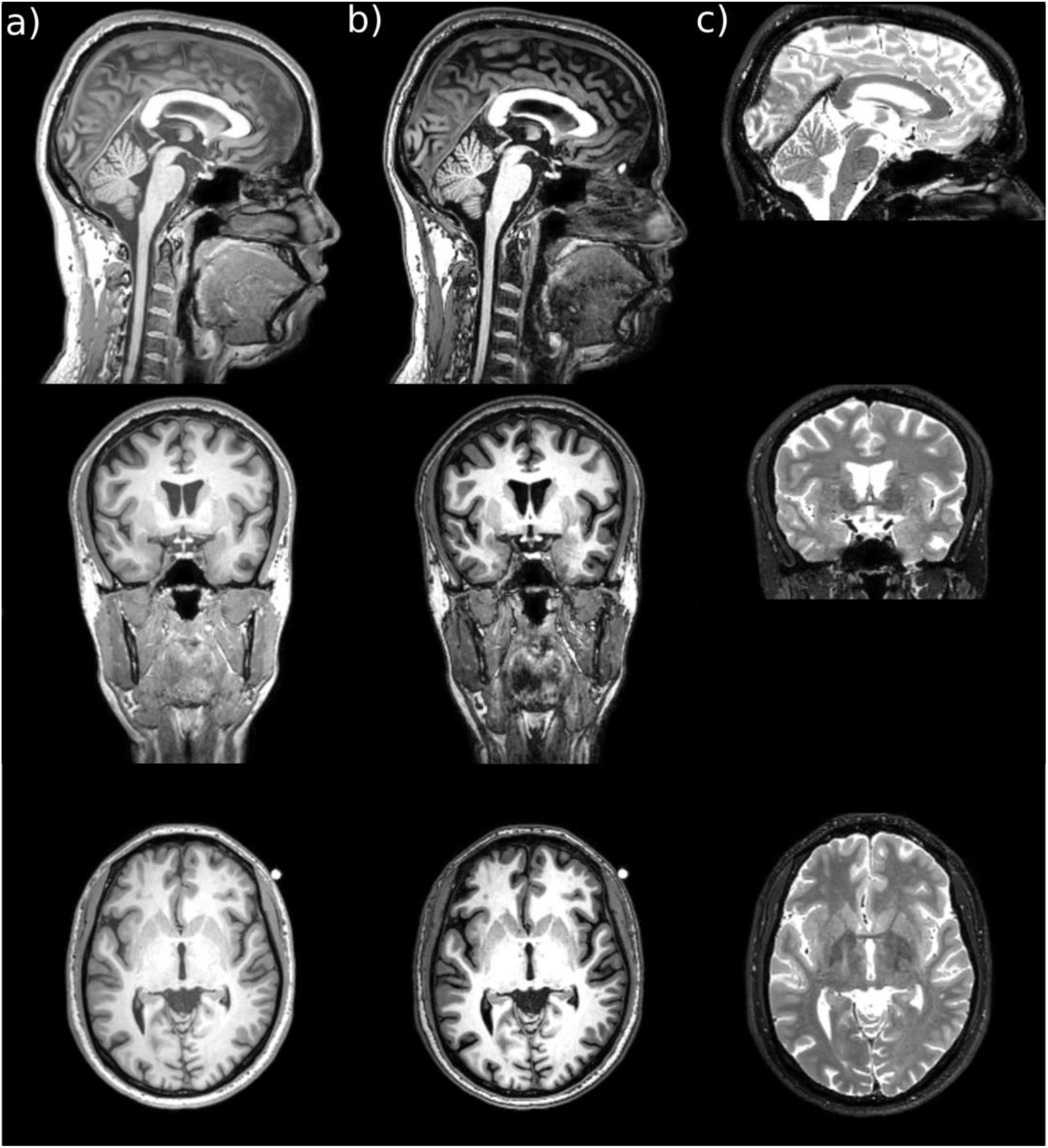
contrast profile of a) T1w, b) T1w-PSIR, and c) T2w scans. Each column shows the image contrast for that image type. The bright dot seen just outside the image in the transverse view of T1w and T1w-PSIR images (lower panel) is the vitamin-E capsule taped to the temple of the subject and is used as a fiducial marker for ease of left and right demarcation. Note that these images are AC-PC aligned.

**Table S1:** Scanning parameters for T1-weighted, T1-weighted PSIR, and T2-weighted scans

| Parameter | T1-weighted | T1-weighted PSIR | T2-weighted |
| --- | --- | --- | --- |
| **FOV (FH × AP × RL) (mm)** | 256 × 256 × 192 | 256 × 256 × 192 | 149.5 × 240 × 176 |
| **Acq orientation** | Sagittal | Sagittal | Transverse |
| **Scan mode / technique** | 3D / FFE | 3D / FFE | MS / SE |
| **Acq voxel size (MPS) (mm)** | 1 × 1 × 1 | 1 × 1 × 1 | 1 × 1.07 × 1 |
| **Recon voxel size (MPS) (mm)** | 1 × 1 × 1 | 1 × 1 × 1 | 1 × 1 × 1 |
| **Acq matrix (MP)** | 256 × 256 | 256 × 256 | 240 × 165 |
| **Recon matrix** | 256 | 256 | 240 |
| **SENSE factor (AP / RL)** | 1 / 2 | 2 / 1.75 | No acceleration |
| **Number of slices** | 192 | 192 | 136 |
| **Slice gap (mm)** | 0 | 0 | 0.1 |
| **Fold-over direction** | AP | AP | RL |
| **Fat-shift direction** | F | F | P |
| **Fast-imaging mode** | TFE | TFE | TSE |
| **Shot mode** | Single-shot | Multishot | Multishot |
| **TFE/TSE factor** | NA | 165 | 15 |
| **TR / TE (ms)** | 6.5 / 2.9 | 7.9 / 3.5 | 3000 / 80 |
| **Flip angle (degrees)** | 9 | 8 | 90 |
| **Scan percentage (%)** | 100 | 100 | 93.75 |
| **Scan duration (mm:ss)** | 05:39.2 | 08:59.2 | 04:48.0 |

**FOV:** field of view; **FH:** foot to head; **AP:** anterior to posterior; **RL:** right to left; **MPS:** measurement, phase, slice; **FFE:** fast field echo; **MS:** multiple slices; **SE:** spin echo; **TFE:** turbo field echo; **TSE:** turbo spin echo; **acq:** acquisition; **recon:** reconstruction

### Functional scans

Critical scan parameters related to acquisition of resting state, and task-based fMRI (TRENDS and VFT) are presented in **Table S2**. Parameters related to reference scans (scans with opposite phase encoding directions: AP and PA) are summarized in **Table S3**.

**Table S2:**
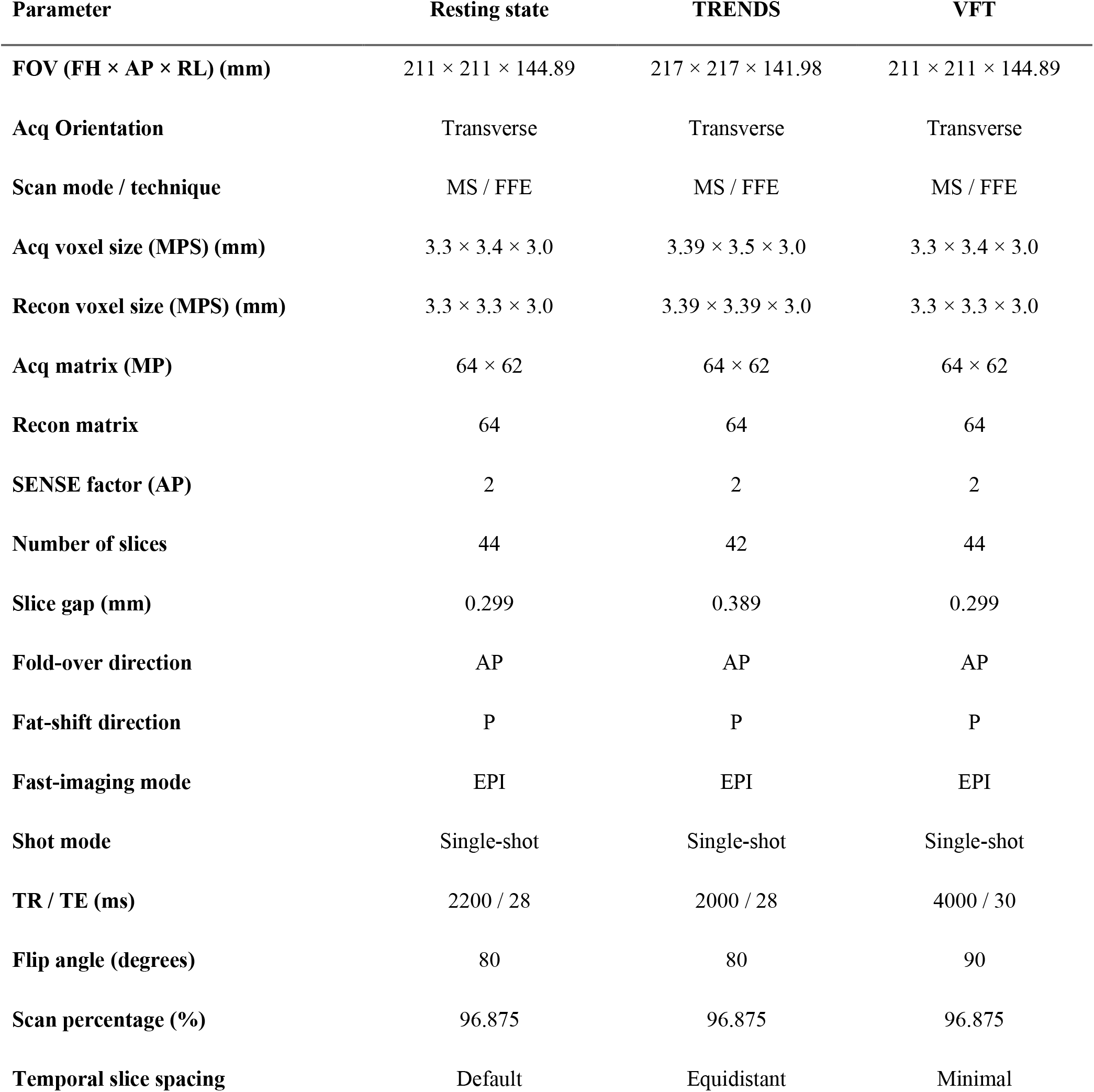
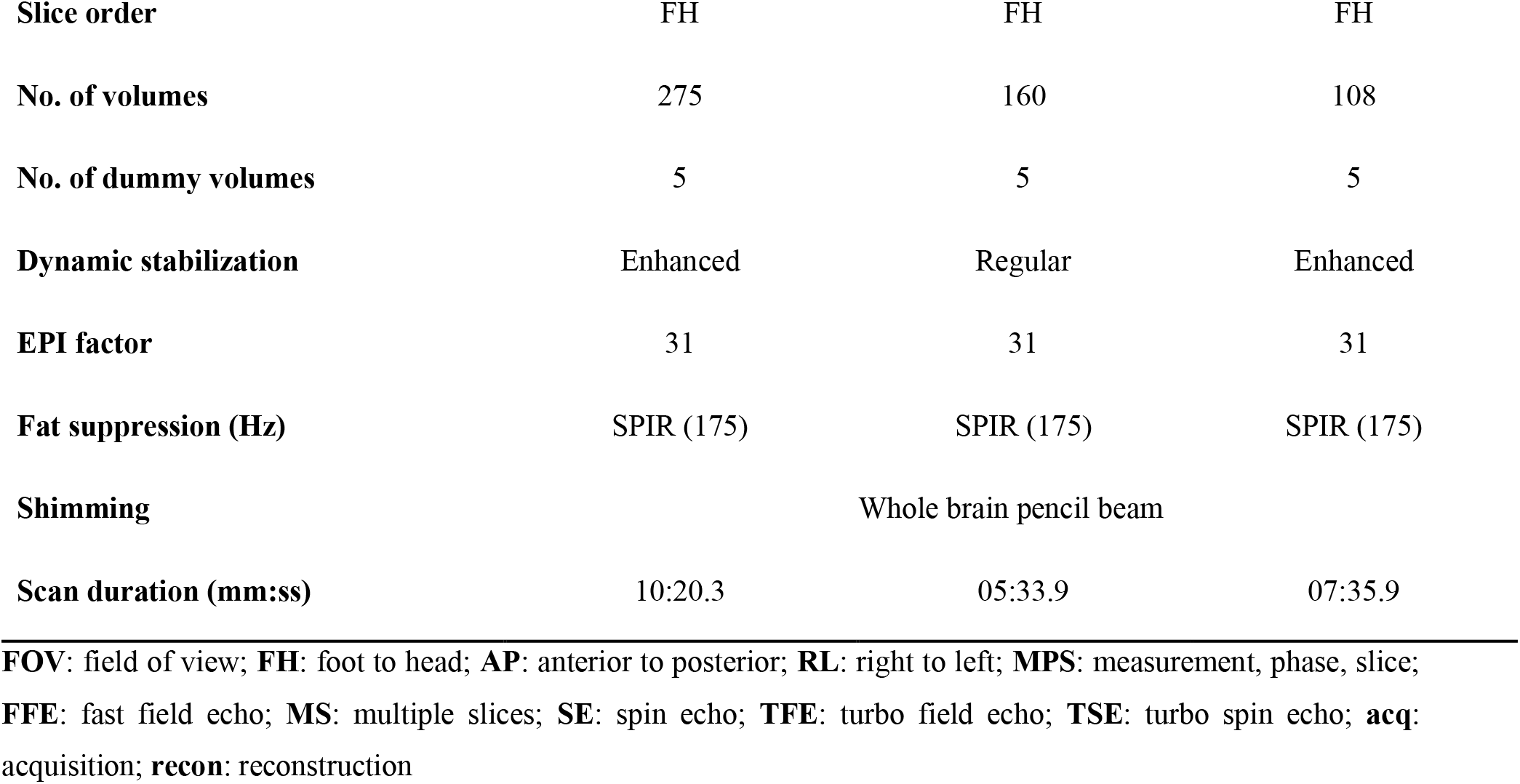
Scanning parameters for resting state fMRI and task-based fMRI: TRENDS and VFT

**Table S3:**
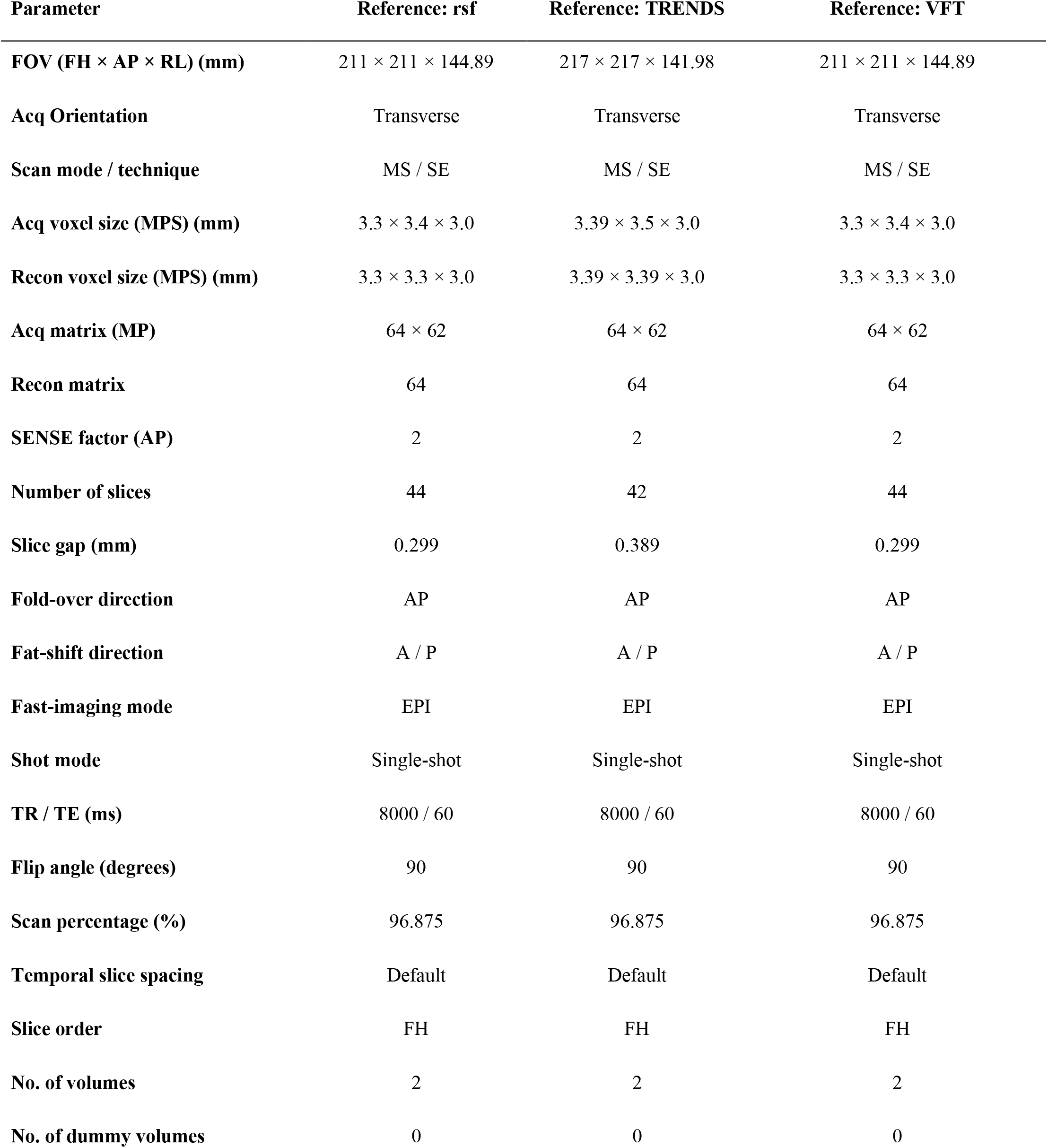
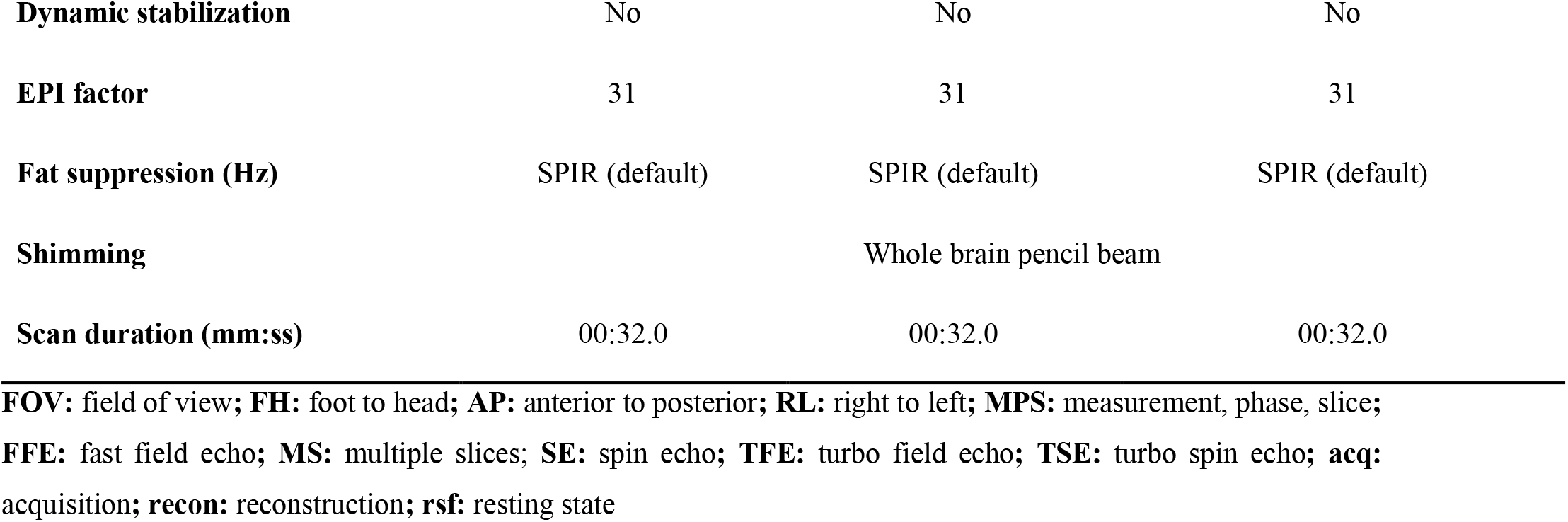
Scanning parameters for reference scan (forward and reverse directions) for resting state fMRI, and task-based fMRI: TRENDS and VFT

### Diffusion scans

A summary of scan parameters for diffusion weighted scans are presented in **Table S4**.

**Table S4:** Scanning parameters for multi-shell diffusion weighted acquisition and reference scan (forward and reverse direction)

| Parameter | Diffusion: multi-shell | Diffusion: reference scan |
| --- | --- | --- |
| **FOV (FH × AP × RL) (mm)** | 240 × 240 × 140 | 240 × 240 × 140 |
| **Acq Orientation** | Transverse | Transverse |
| **Scan mode / technique** | MS / SE | MS / SE |
| **Acq voxel size (MPS) (mm)** | 2.50 × 2.55 × 2.50 | 2.50 × 2.55 × 2.50 |
| **Recon voxel size (MPS) (mm)** | 2.50 × 2.50 × 2.50 | 2.50 × 2.50 × 2.50 |
| **Acq matrix (MP)** | 96 × 94 | 96 × 94 |
| **Recon matrix** | 96 | 96 |
| **SENSE factor (AP)** | 2 | 2 |
| **Number of slices** | 56 | 56 |
| **Slice gap (mm)** | 0 | 0 |
| **Fold-over direction** | AP | AP |
| **Fat-shift direction** | P | A / P |
| **Fast-imaging mode** | EPI | EPI |
| **Shot mode** | Single-shot | Single-shot |
| **TR / TE (ms)** | 7221 / 100 | 8000 / 60 |
| **Flip angle (degrees)** | 90 | 90 |
| **Scan percentage (%)** | 97.92 | 97.92 |
| **Temporal slice spacing** | Default | Default |
| **Slice order** | Default | Default |
| **No. of volumes** | 80 | 2 |
| **EPI factor** | 47 | 47 |
| **Shimming** | Whole brain pencil beam | Whole brain pencil beam |
| **Fat suppression (Hz)** | SPIR (Default) | SPIR (Default) |
| **Scan duration (mm:ss)** | 10:35.4 | 00:32.0 |

**FOV:** field of view; **FH:** foot to head; **A:** anterior; **P:** posterior; **AP:** anterior to posterior; **RL:** right to left; **MPS:** measurement, phase, slice; **acq:** acquisition; **recon:** reconstruction; **EPI:** echo planar imaging

## Brief description of task fMRI

### TRENDS

Tool for Recognition of Emotions in Neuropsychiatric Disorders (TRENDS) is a task that includes a presentation of still pictures showing five basic emotions of sadness, fear, anger, surprise, and disgust at high and low intensities along with neutral expressions in a block design. All the images corresponding to each emotion are randomized. Participants are instructed to lie down and keep their eyes open to view the images appearing on the screen. More details of TRENDS can be found in (Behere et al., 2008)

### Verbal fluency task (VFT)

The verbal fluency task consists of two conditions: word repetition (WR) and word generation (WG). During the WR condition, participants are shown the word “pass” on the screen and instructed to say “pass” as soon as possible. In the WG condition, participants are shown the name of a category on the screen and are expected to generate a unique example of that category as soon as possible. Each block consists of seven trials. Participants are instructed to not repeat the same example and to say “pass” in case they are unable to come up with a new example word. This is done to ensure same number of vocal responses across blocks. The responses are picked up by the MR compatible microphone described previously. The acquisition is clustered such that the temporal spacing between slices in a volume is minimum, thereby giving a “silent period” during which participants vocalize the generated word. More details of VFT can be found in (John, Halahalli, Vasudev, Jayakumar, & Jain, 2011)

## Post-acquisition protocol

All acquired data in Digital Imaging and Communications in Medicine (DICOM) format are screened using a batch version of DicomBrowser (Archie & Marcus, 2012) to detect any possible data entry errors. We check (and correct, if needed) the following DICOM tags to ensure that the records are correct: participant name, age, gender, date of birth, hospital ID, and internal reference numbers.

Once the integrity of the data is established, we use Chris Rorden’s dcm2niix (https://github.com/rordenlab/dcm2niix) software to convert DICOM format data to Neuroimaging Informatics Technology Initiative (NIfTI) format. Philips DICOM data has two types of intensity scaling values: Philips precise and display values; based on the suggestions in (Chenevert et al., 2014), we explicitly use Philips precise values when converting the data to NIfTI. To facilitate the process of reorganizing the NIfTI dataset to the Brain Imaging Data Structure (BIDS) (Gorgolewski et al., 2016), we rename the participant folders in the following format: “sub-YYYYMMDD9digitID” where YYYYMMDD refers to the date of MRI acquisition and the 9 digit ID is an internal reference ID that can be used to track additional information about this subject. The first three digits of this ID pertains to the visit number of the subject (for example 101 for the first visit, 102 for the second visit); the next six digits are the unique participant identifier. Additional JavaScript Object Notation (JSON) sidecar files are automatically generated when converting DICOM data to NIfTI. At this stage, we pass the subjects’ NIfTI folder through a MATLAB program that renames and reorganizes the NIfTI files so that the data becomes BIDS compliant.

All acquired DICOM data are anonymized to remove any identifiable information of the subject. Specifically, we remove the participant name, age, gender, date of birth, all fields containing hospital IDs, and internal reference numbers. Once the data has been converted to NIfTI format, we additionally subject each structural image to a de-facing protocol by using the de-facing routine of SPM12 (https://www.fil.ion.ucl.ac.uk/spm/).

## Phantom Positioning

Both the agar gel phantom and then HPD phantom are placed in the 32-channel head coil; care is taken to ensure that the placement and planning of acquisition is symmetrical and as similar as possible to the human acquisition. **Figure S2** shows the placement of then phantom in the head coil while **Figure S3** shows the planning of the acquisition.

**Figure S2:**
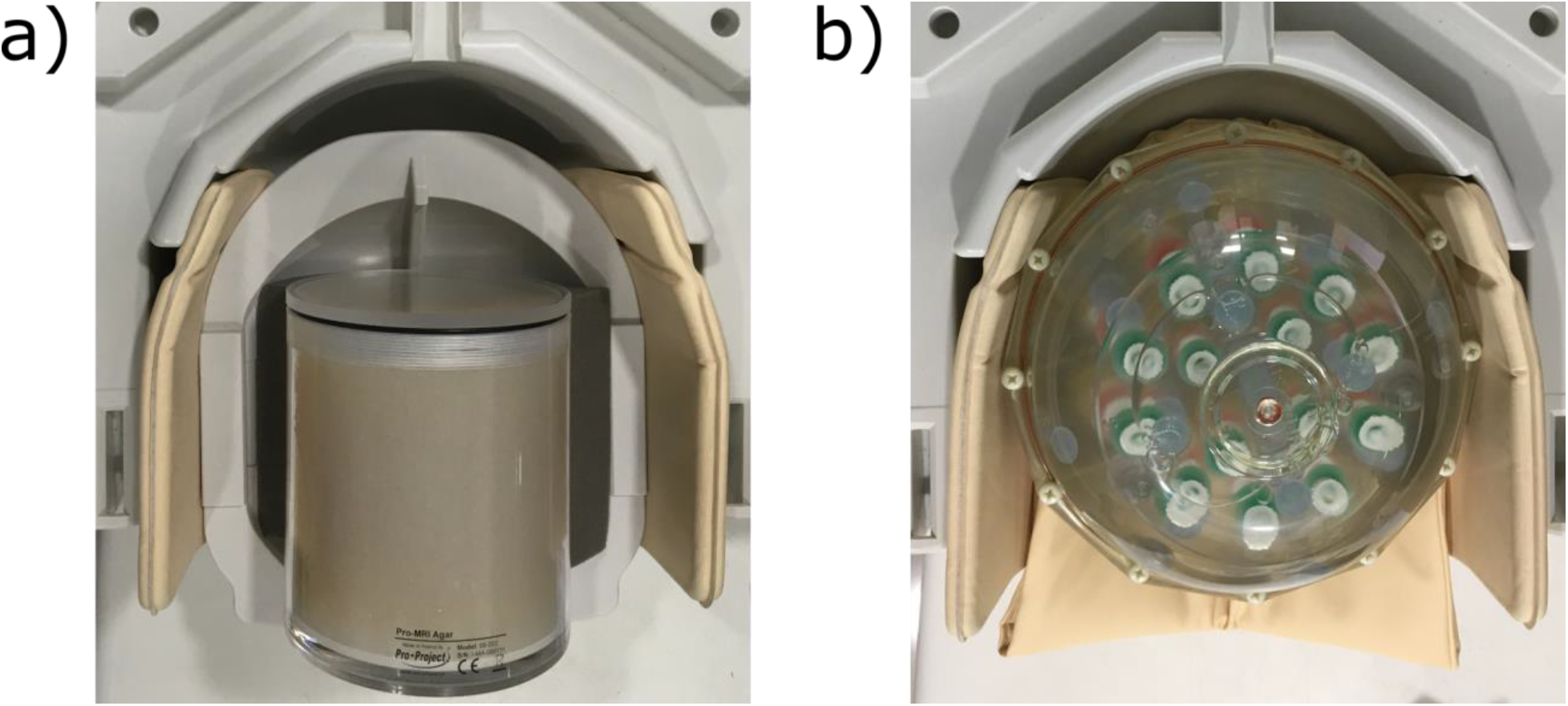
Placement of the a) gel and b) HPD phantom in the 32-channel head coil

**Figure S3:**
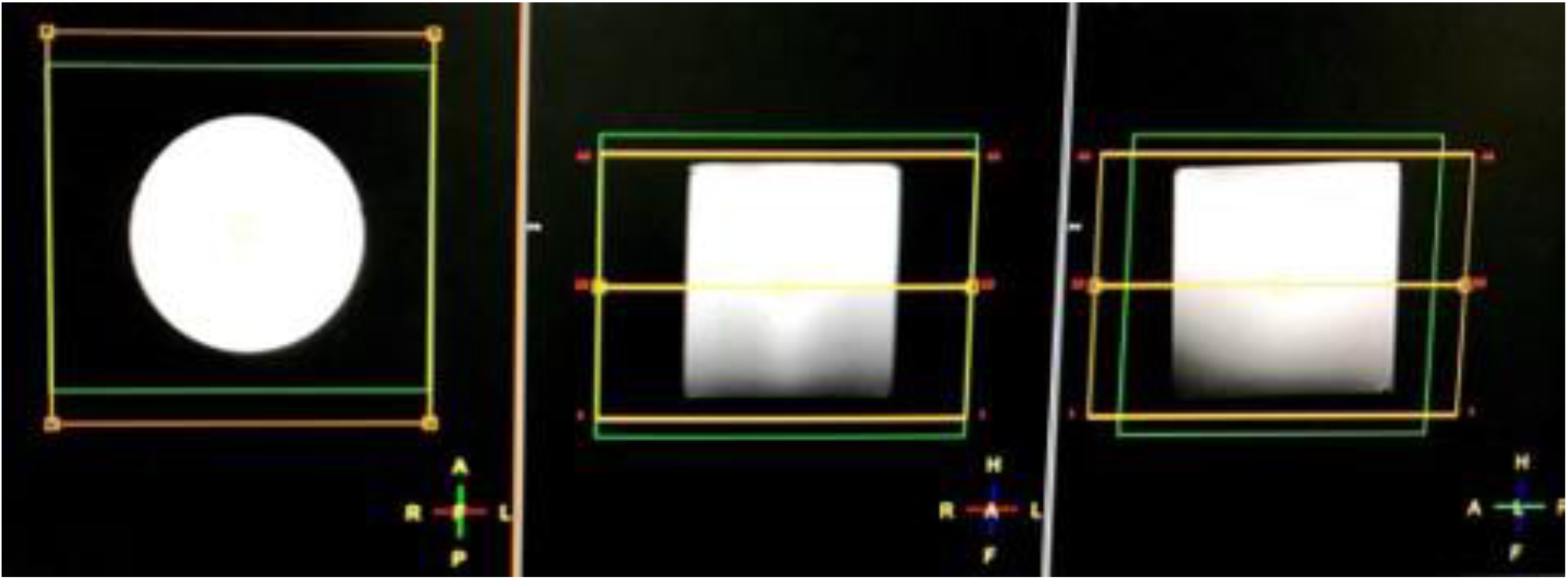
Planning of gel agar phantom acquisition; the yellow box shows the field of view while the green box shows the planning of pencil beam shimming method in the three orientations

## Analysis of gel agar phantom

For the analysis of gel agar phantom data, we adapted the code from LAB-QA2GO (https://github.com/vogelbac/LAB-QA2GO). Foremost, since our acquisition did not include dummy volumes, we start by ignoring the first three volumes. For our acquisitions, the field of view was the same as the one used for human acquisition. Therefore, the images include the entire phantom (including background on top and bottom). However, the slices covering the head and tail part of the phantom would merely include the casing of the phantom and not actual gel. Considering these slices for analysis would be erroneous. Further, since the first step in the analysis method is to create a binary mask using Otsu’s method, these slices would result in a fragmented binary mask which would be problematic for all further calculations.

To account for these slices, we calculate the number of connected components for each slice for each volume in the binary mask image. Any slice where the mask is “fragmented” (due to edge effects), the number of connected components would be more than one. We detect the beginning and ending of the phantom in this manner. For our phantom, we need to extend these detected slices by a few more to reach the part of the phantom where gel exists (see **Error! Reference source not found.Error! Reference source not found.**). We have found that adding two more slices on each side (after detecting the beginning and the ending of then phantom) usually gives satisfactory results. This step is likely specific to our phantom. For these slices, we replace the calculated binary mask with the mask from the center slice. We ignore these slices when calculating percent signal ghosting and percent signal change.

On comparing our phantom data with the sample data from LAB-QA2GO, we noticed that our phantom was larger in diameter. This can interfere with the ‘ghosting detection’ module which attempts to place a 10×10 phantom on each side of the phantom (in the background), as there might not be enough space. We noticed that on the left and right side, depending on the binary mask, the ROI placement would often fail. To compensate for this, we iteratively reduce the size of the ROI till an ROI placement can be achieved. For our scans, the smallest ROI size needed was 9, though a size of 10 worked for some of the cases.

**Figure S4:**
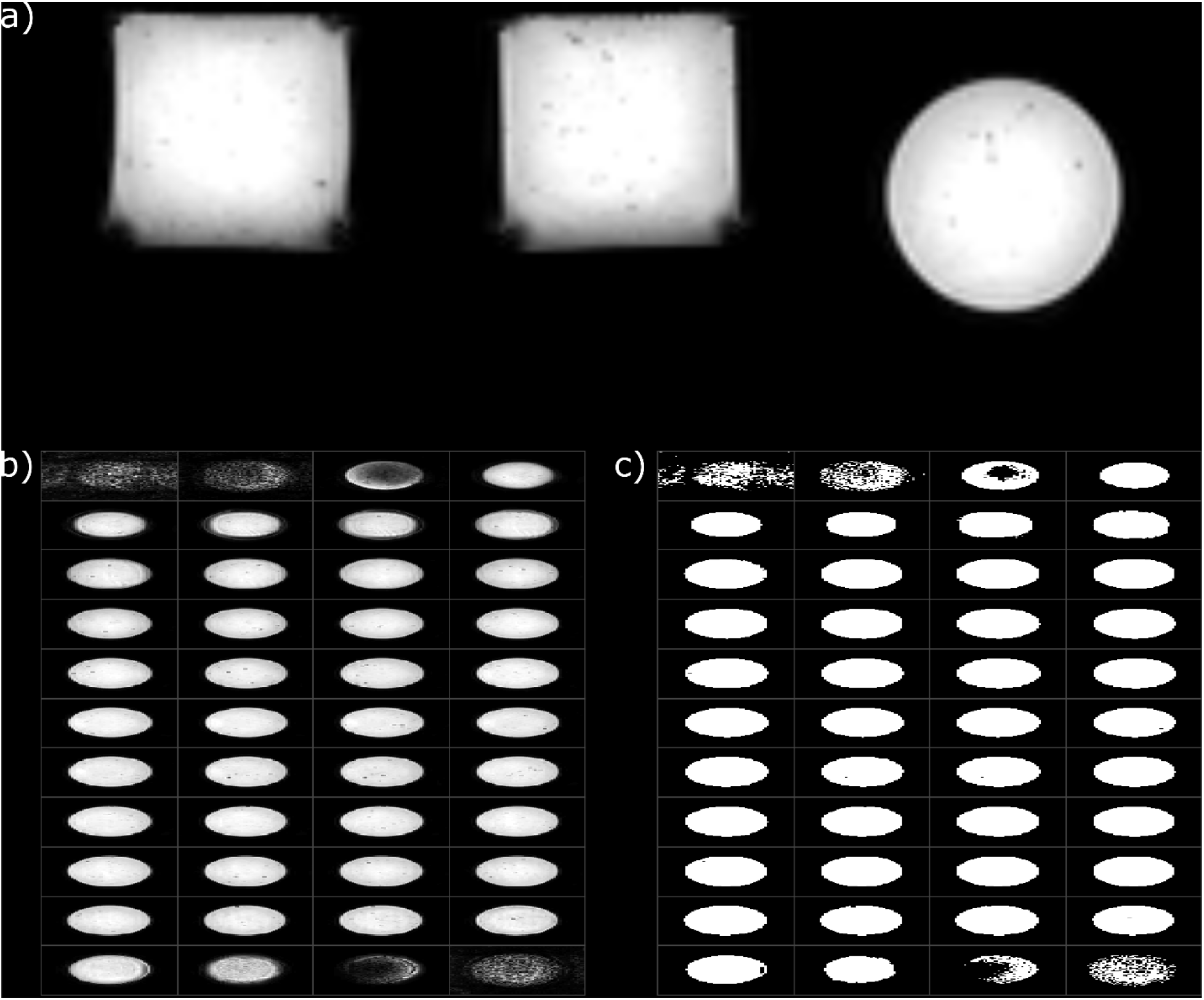
a) Example slice in all three planes for gel agar phantom data; b) all slices for a volume in the transverse plane; c) binary mask created using Ostu’s method for the same slices as in b); notice the edge effects in the first few slices on the top and bottom of the image in a). These slices are the beginning and end of the phantom where gel agar is not present; these slices would result in a poor binary mask (as seen in c)) and would interfere with all calculation steps. We detect these slices and replace their binary mask with the mask from the middle slice. We also ignore these slices when calculating percent signal ghosting and percent signal change

## Scan schedule: agar gel phantom

**Table S5:** Breakup of the number of agar gel phantoms acquisitions per month

| Month of acquisition | Number of scans |
| --- | --- |
| 2018-Apr | 03 |
| 2018-May | 02 |
| 2018-Jun | 04 |
| 2018-Jul | 04 |
| 2018-Aug | 05 |
| 2018-Sep | 03 |
| 2018-Oct | 04 |
| 2018-Nov | 04 |
| 2018-Dec | 03 |
| 2019-Jan | 04 |
| 2019-Feb | 04 |
| 2019-Mar | 05 |
| 2019-Apr | 03 |
| 2019-May | 04 |
| 2019-Jun | 04 |
| 2019-Jul | 03 |
| 2019-Aug | 24 |
| 2019-Sep | 21 |
| 2019-Oct | 23 |
| 2019-Nov | 22 |
| 2019-Dec | 23 |

## Scan schedule: HPD phantom (proton density)

**Table S6:** Breakup of the number of HPD phantom proton density acquisitions per month

| Month of acquisition | Number of scans |
| --- | --- |
| 2018-Apr | 01 |
| 2018-May | 03 |
| 2018-Jun | 03 |
| 2018-Jul | 04 |
| 2018-Aug | 03 |
| 2018-Sep | 04 |
| 2018-Oct | 03 |
| 2018-Nov | 01 |
| 2018-Dec | 00 |
| 2019-Jan | 00 |
| 2019-Feb | 01 |
| 2019-Mar | 01 |
| 2019-Apr | 01 |
| 2019-May | 01 |
| 2019-Jun | 01 |
| 2019-Jul | 01 |
| 2019-Aug | 04 |
| 2019-Sep | 04 |
| 2019-Oct | 04 |
| 2019-Nov | 03 |
| 2019-Dec | 05 |

## Scan schedule: Reliability analysis

**Table S7:** Date and approximate time of acquisition for T1-weighted and diffusion weighted scans for volunteer test-retest reliability scans

| Date of acquisition | Time of T1-weighted scan | Time of DWI scan |
| --- | --- | --- |
| 23-July-2019 | 15:14:26 | 16:02:41 |
| 24-July-2019 | 14:50:46 | 15:36:41 |
| 27-July-2019 | 17:09:08 | 17:54:19 |
| 19-Aug-2019 | 14:22:07 | 15:07:07 |
| 20-Aug-2019 | 14:39:41 | 15:24:37 |
| 21-Aug-2019 | 14:18:33 | 15:05:48 |
| 26-Sep-2019 | 14:30:14 | 15:18:17 |
| 27-Sep-2019 | 15:30:45 | 16:18:02 |
| 30-Sep-2019 | 15:23:59 | 16:11:26 |
| 17-Oct-2019 | 16:06:43 | 16:20:14 |
| 18-Oct-2019 | 16:23:14 | 17:09:25 |
| 19-Oct-2019 | 15:41:49 | 16:15:44 |
| 14-Nov-2019 | 14:33:37 | 14:46:13 |
| 15-Nov-2019 | 14:37:39 | 14:50:21 |
| 16-Nov-2019 | 15:34:12 | 15:46:45 |
| 17-Dec-2019 | 14:51:27 | 15:24:16 |
| 18-Dec-2019 | 14:35:23 | 15:03:17 |
| 19-Dec-2019 | 14:32:52 | 14:51:22 |

## Medication details during reliability analysis

**Table S8:** Details of medication for volunteer for test-retest reliability scans

| Date of acquisition | Medication details |
| --- | --- |
| 23<sup>rd</sup> July 2019 | Pregabalin 75mg at 11:25 AM (+Nifedipine 10mg SR at night-time) |
| 24<sup>th</sup> July 2019 | Pregabalin 75mg at 11:33 AM (+Nifedipine 10mg SR at night-time) |
| 27<sup>th</sup> July 2019 | Pregabalin 75mg at 10:35 AM (+Nifedipine 10mg SR at night-time) |
| 19<sup>th</sup> August 2019 | Pregabalin 75mg at 12:30 PM |
| 20<sup>th</sup> August 2019 | Pregabalin 75mg at 09:00 AM |
| 21<sup>st</sup> August 2019 | Pregabalin 75mg at 08:30 AM |

## Quality check measures example plots

### DVARS calculation

**Figure S5:**
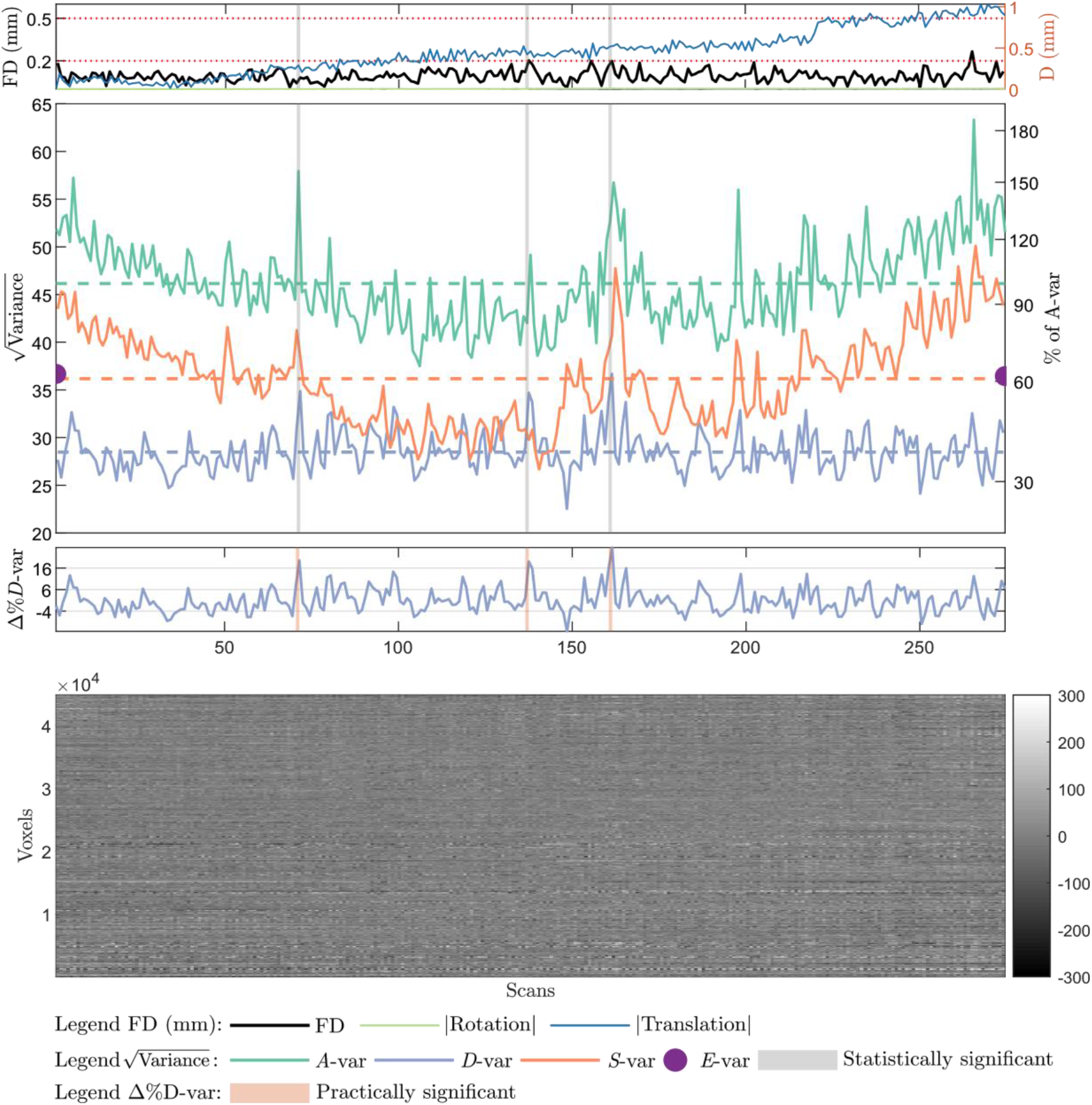
Output of motion detection using the modified DVARS implementation (Afyouni and Nichols, 2018). The top panel of the figure shows head movement of a given subject as translations and rotations, along with framewise displacement calculated as sum of absolute values of the derivations of the six realignment parameters (three translations and three rotations); the second panel from the top shows sum of squares decomposition of data into *A-*var (total sum of squares at a time point t) and *D*-var, *S*-var, and *E*-var corresponding to fast, slow, and edge variability terms; pairs of time points which are detected as outliers (using a statistical threshold of *α*<0.05 corrected for multiple comparison using a Bonferroni approach) are marked in gray; the third panel from the top depicts the percent change in *D*-var term with outliers (detected using a liberal practical threshold of 5% *D*-var change) marked in orange colour; the lower panel of the figure depicts a carpet plot showing fMRI signal from all voxels over time; this is helpful in assessing any major noise in the data. Note that a scaling of 1/100 has been applied to the data.

### Signal and brainmask profile

**Figure S6:**
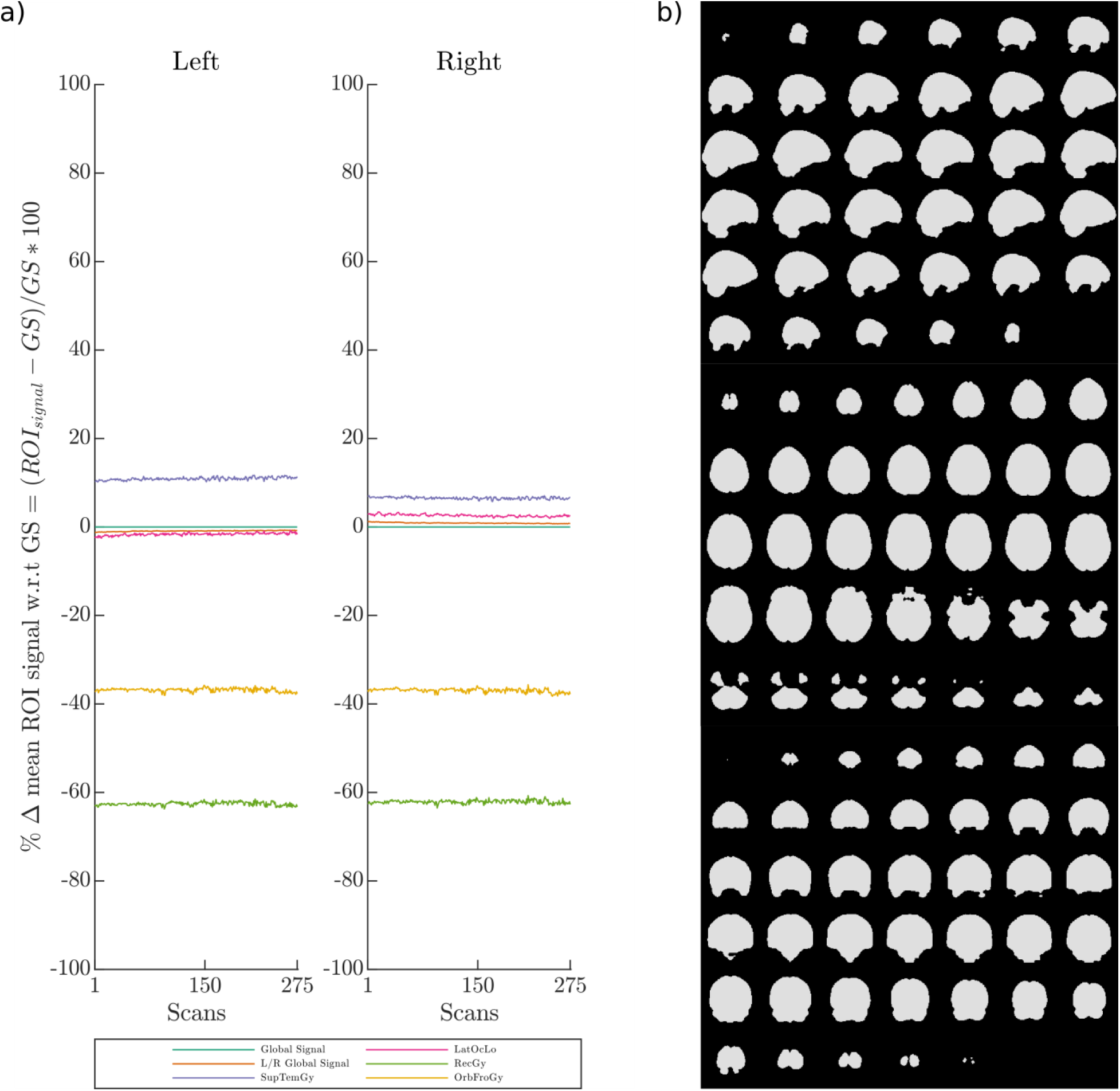
a) Percentage change in mean ROI signal with respect to global signal (GS) across different regions which may or may not be susceptible to signal loss; b) An example of brainmask generated by preserving only voxels which have a mean signal of more than 80% of global signal. These brainmask profiles can be used to eliminate subjects where a large number of voxels have poor signal.

### MRI screening form

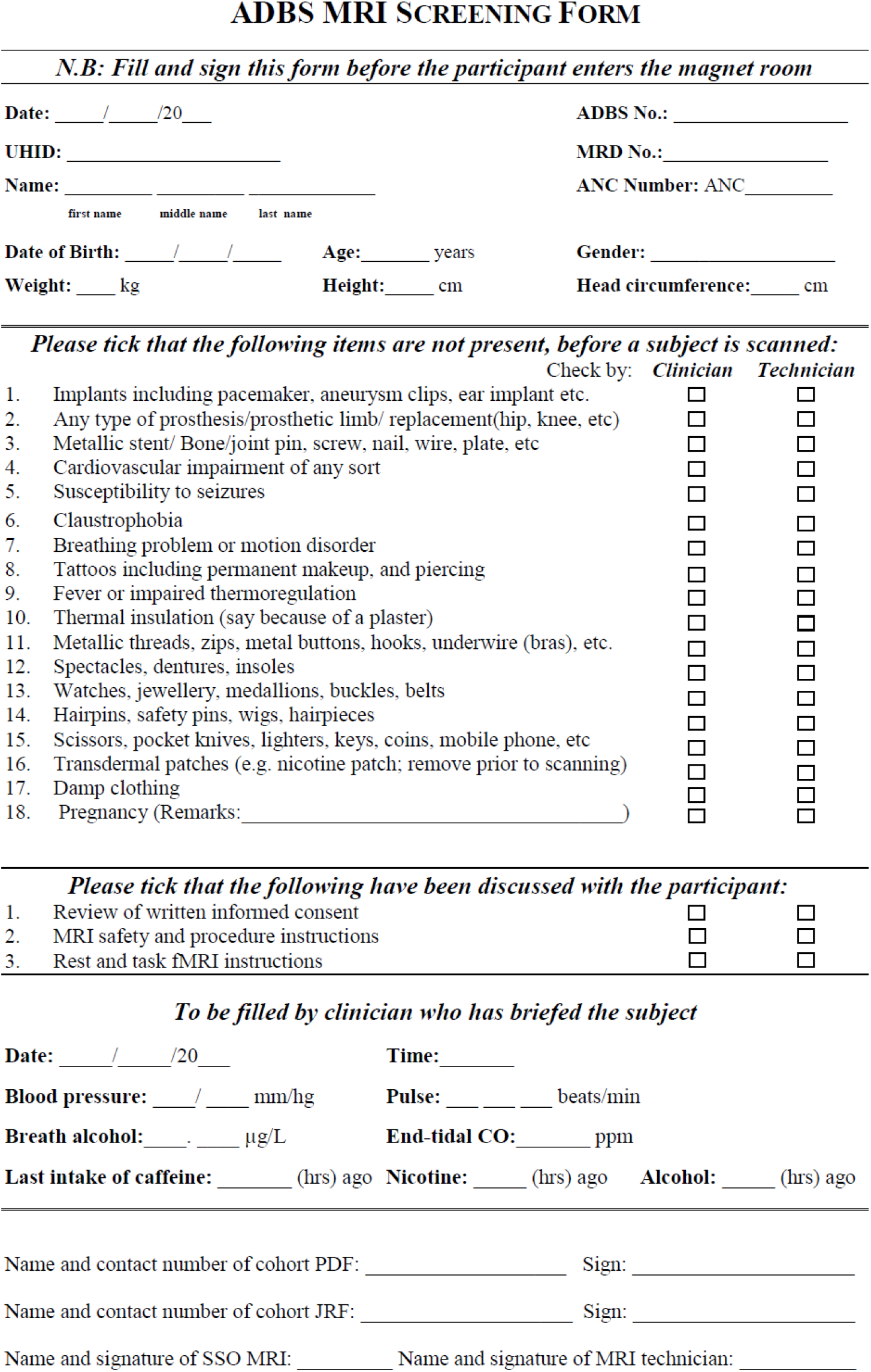

|  | Check by: | Clinician | Technician |
| --- | --- | --- | --- |
| 1. Implants including pacemaker, aneurysm clips, ear implant etc. |  | ☐ | ☐ |
| 2. Any type of prosthesis/prosthetic limb/ replacement(hip, knee, etc) |  | ☐ | ☐ |
| 3. Metallic stent/ Bone/joint pin, screw, nail, wire, plate, etc |  | ☐ | ☐ |
| 4. Cardiovascular impairment of any sort |  | ☐ | ☐ |
| 5. Susceptibility to seizures |  | ☐ | ☐ |
| 6. Claustrophobia |  | ☐ | ☐ |
| 7. Breathing problem or motion disorder |  | ☐ | ☐ |
| 8. Tattoos including permanent makeup, and piercing |  | ☐ | ☐ |
| 9. Fever or impaired thermoregulation |  | ☐ | ☐ |
| 10. Thermal insulation (say because of a plaster) |  | ☐ | ☐ |
| 11. Metallic threads, zips, metal buttons, hooks, underwire (bras), etc. |  | ☐ | ☐ |
| 12. Spectacles, dentures, insoles |  | ☐ | ☐ |
| 13. Watches, jewellery, medallions, buckles, belts |  | ☐ | ☐ |
| 14. Hairpins, safety pins, wigs, hairpieces |  | ☐ | ☐ |
| 15. Scissors, pocket knives, lighters, keys, coins, mobile phone, etc |  | ☐ | ☐ |
| 16. Transdermal patches (e.g. nicotine patch; remove prior to scanning) |  | ☐ | ☐ |
| 17. Damp clothing |  | ☐ | ☐ |
| 18. Pregnancy (Remarks: _____) |  | ☐ | ☐ |

*Please tick that the following have been discussed with the participant:*
|  |  |  |
| --- | --- | --- |
| 1. Review of written informed consent | ☐ | ☐ |
| 2. MRI safety and procedure instructions | ☐ | ☐ |
| 3. Rest and task fMRI instructions | ☐ | ☐ |

*To be filled by clinician who has briefed the subject*
Date: \_\_\_\_/\_\_\_\_/20\_\_\_\_ Time: \_\_\_\_\_ Blood pressure: \_\_\_\_/\_\_\_\_ mm/hg Pulse: \_\_\_\_ beats/min Breath alcohol: \_\_\_\_ µg/L End-tidal CO: \_\_\_\_ ppm Last intake of caffeine: \_\_\_\_ (hrs) ago Nicotine: \_\_\_\_ (hrs) ago Alcohol: \_\_\_\_ (hrs) ago
Name and contact number of cohort PDF: \_\_\_\_\_ Sign: \_\_\_\_\_
Name and contact number of cohort JRF: \_\_\_\_\_ Sign: \_\_\_\_\_
Name and signature of SSO MRI: \_\_\_\_\_ Name and signature of MRI technician: \_\_\_\_\_

